# Cortico-subcortical multi-head self-attention as a substrate for cognitive performance

**DOI:** 10.64898/2026.08.06.743205

**Authors:** Vasanth Kumar Babu, Arno Granier, Timothée Proix, Titas Balvocius, Marmaduke Woodman, Ausra Saudargiene, Viktor Jirsa, Walter Senn

## Abstract

The neocortex is central to mammalian cognition, yet a computational framework that is both biologically constrained and capable of complex cognitive tasks remains missing. Here we show that cortico-thalamic circuits are well suited to implement multi-head self- and cross-attention, the mechanism behind the cognitive abilities of transformer networks. We propose that layer 2/3 pyramidal cells maintain a recurrent key-value memory, which layer 5 pyramidal cells read out in response to a query. Keys, values and queries map onto core and matrix thalamo-cortical projections, distributed across the micro- and macro-columns of a cortical area, and each cortical area forms one attention head. The same microcircuit also computes the prediction errors that drive synaptic plasticity, while a reward-prediction error from the basal ganglia gates the cortical output and the recall of hippocampal memories. The trained network aligns with human intracranial recordings during speech perception. Our cortico-subcortical attention circuit thus offers a biologically grounded substrate for mammalian cognition.

## Introduction

The neocortex is central to the cognitive abilities of the mammalian brain, from perceiving complex scenes to learning and producing language, and reasoning over abstract relationships. Traditional computational models of the cortex in neuroscience have struggled to translate anatomical detail into cognitive capability: they capture local circuit dynamics or single-task performance, but fail to match the breadth and depth of cortical function. Connectome-constrained brain network models have succeeded in reproducing the brain activity of individual subjects [1, 2, 3], but remain limited to functionally inert brain dynamics such as seizures, resting-state activity and responses to brain stimulation. A computational framework that is both constrained by biology and powerful enough to perform hard cognitive tasks remains missing.

Recent advances in artificial intelligence (AI), driven most prominently by transformer networks [4], produced AI models that match or exceed human performance across a growing range of cognitive tasks. The high performance of transformer networks is not limited to natural language processing, but extends to any domain where inputs can be expressed as sequences of ‘tokens’, including abstract reasoning [5] and vision [6], where a token encodes a word or a foveal image patch, for instance. At the heart of transformer networks lies the self-attention mechanism, a scalable and parallelizable computation that captures long-range dependencies within a sequence. Identifying a cortical implementation of self-attention would therefore offer a candidate mechanism for how the cortex achieves its cognitive abilities.

The neocortex of mammals is built of canonical computational motifs [7, 8, 9], repeated both in parallel and in series [10, 11]. Concerted efforts have provided cell-type-specific data on the local components of this canonical cortical circuit [12, 13, 14]. The brain also shows a hierarchical organization, developed through evolution, with sub-cortical structures, such as basal ganglia and the midbrain, calculating values and reward-prediction errors [15, 16]. Memories are likewise hierarchically organized, with hippocampal episodic memory providing contextual information to semantic cortical memories [17, 18]. There is also growing computational interest in the interaction between the cerebral cortex and the thalamus, and in its contribution to learning [19], perception [20] and awareness [21, 22]. We are finally beginning to understand the involvement of dendrites in error representation during learning and behavior [23, 24].

Previous work on biologically plausible neural implementations of self-attention has focused on a single head, aiming to model either the hippocampal formation [25], the role of glial cells [26], or general mappings to neuroscience [27, 28]. Biological plausibility has also been argued from the equivalence between self-attention and the update in modern continuous Hopfield networks [29, 30]. On the other hand, error backpropagation, the workhorse of modern AI, was proposed to be implemented in local cortical microcircuits [31, 32], and the minimization of dendritic prediction errors was suggested as a core element connecting cellular features with general computation, e.g. formulated in the neuronal least-action principle [33, 34].

Here, we propose that cortico-thalamic circuits, including their cell types, pathways and interactions, are well suited to implement a computation akin to multi-head self- and cross-attention, and we present gradient-descent plasticity rules that could operate in the brain. We build on the formulation of linear self-attention as a recurrent key-value memory system [35, 36, 37]. We show how cortical layer 2/3 (L2/3) pyramidal neurons could form a recurrent key-value memory integrating thalamic inputs, while layer 5 (L5) pyramidal neurons could perform retrieval by the current query. Crucially, we highlight how brain-specific structures, such as the extensively branched cortical connectivity patterns, the thalamic gateway to cortex, the reward-based modulation of cortical activity, and the hippocampal memory extension, improve cognitive performance. Our quest for a cortical implementation of self-attention further reveals an anatomical duality between representing attention during inference and representing prediction errors during learning.

The structural analogies we report are meant as a guide: they suggest a mechanistic understanding of cognition in terms of key-value memories and attention, and at the same time point to brain features that have no direct counterpart in artificial transformers. In our cortical-subcortical model, for example, the L2/3 key-value memory is a working memory into which hippocampal episodic memories are recalled, and the basal ganglia use a reward-prediction error to gate both this recall and the cortical output.

The paper is organized as follows: after presenting the theory of key-value memories and self-attention, together with the mapping to thalamo-cortical circuits, we show how these circuits can robustly learn small-scale natural language generation. We then generalize the concept of self- and cross-attention to inter-areal cortical interactions across sensory-motor streams, and show how these cortical features make the language task robust against stochastic variations in the cortical connectivity. We further map the network activity onto intracortical activity recorded in human participants during speech perception, showing the alignment of model and data. We next show how dendritic gain modulation by the unsigned reward-prediction error sharpens the cortical representation for translation during multi-language exposure. Finally, we explain how task-specific hippocampal memories can be recalled into the cortical memory, and how the same thalamo-cortical microcircuits can represent prediction errors required for gradient-based learning. Early ideas of this work were introduced as a preprint [27].

## Results

### Multi-head self-attention for selective memory retrieval

We first present in a nutshell the theory of multi-head self-attention before mapping it to the brain. An attention-head will then be implemented by a cortical area, and the attention mechanism by the cortical layers. In response to an input token ***x***_*t*_ ∈ ℝ^*d*^ (a column vector) at time *t*, heads with index *h* compute keys 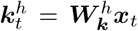 with matrices 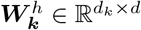 for each head, and values 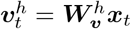 with 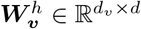. Head-specific memory matrices 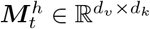 recurrently integrate an association between a new key-value pair, alongside those of past key-value memories. At each discrete time *t*, memory matrices are updated by the outer product of values and keys, while discounting the previous memory,

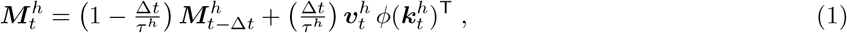

with head-specific time constants *τ* ^*h*^ ≥ Δ*t* and a fixed unit time step Δ*t*. When we model speech processing, we choose a time step of Δ*t* = 50 −100 ms [38], with an integration time constant in the range of seconds, *τ* ^*h*^ = 1 − 2 s. When we are instead interested in the time-continuous processing of sensory inputs, the time step can be made arbitrarily small (say 1 ms). The dendritic transfer function *ϕ* is monotonically increasing (e.g. a threshold-linear function) and applied elementwise. All quantities, except the time constant, are unitless for simplicity.

Upon a new input token ***x***_*t*_, each head (area) constructs a query 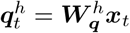 via matrix 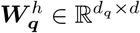, and with this query retrieves a weighted sum of old values from its memory 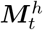. Intuitively, a query is compared with all the stored keys, and proportionally to the overlap between the query and the keys, adds the corresponding values to the output. Formally, the output from head *h* at time *t* is the multiplication of the memory matrix with the query vector,

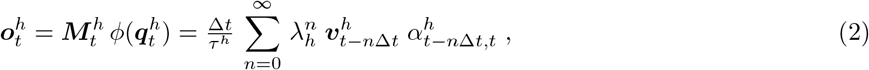

with attention factor 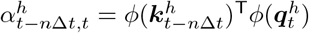 that represents the overlap (scalar product) between the query 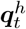 at the current time *t* and the key 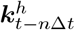 at *n* time steps back (keys and queries have equal dimensions, *d*_*k*_ = *d*_*q*_). The temporal discount factor per Δ*t* is 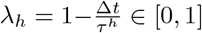. Hence, according to eq. 2, the output from head *h* sums up past value vectors 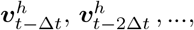 discounted by powers of *λ*_*h*_ and weighted by the attention factor 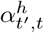.

A context-aware output token ***y***_*t*_ ∈ ℝ^*d*^ is then constructed by summing the retrieved memories 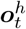 across heads with head-specific output weights 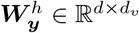,

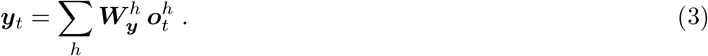

For a discount factor *λ*_*h*_ = 1, eqs. 2 and 3 describe a multi-head, linear self-attention layer with finite, but non-decaying memory trace [36] (see section S2). For simplicity, we consider linear self-attention that does not take into account a soft winner-takes-all mechanism to calculate the attention factors.

### Key-value memories in cortical layer 2/3 pyramidal neurons

L2/3 pyramidal neurons are natural candidates to compute key-value memory 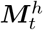 (eq. 1); they form a recurrent network within a cortical column [39, 40, 41] and maintain activity on timescales consistent with short-term memory [42, 43]. We propose that a 2-dimensional cortical sheet of *d*_*v*_ × *d*_*k*_ ensembles of L2/3 pyramidal neurons encodes 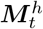 (in this section, we focus on a single head *h* specifying the cortical area). Each ensemble of L2/3 pyramidal neurons is a tightly coupled group of pyramidal neurons representing a single scalar component of 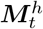 (fig. 1 a).

**Figure 1:**
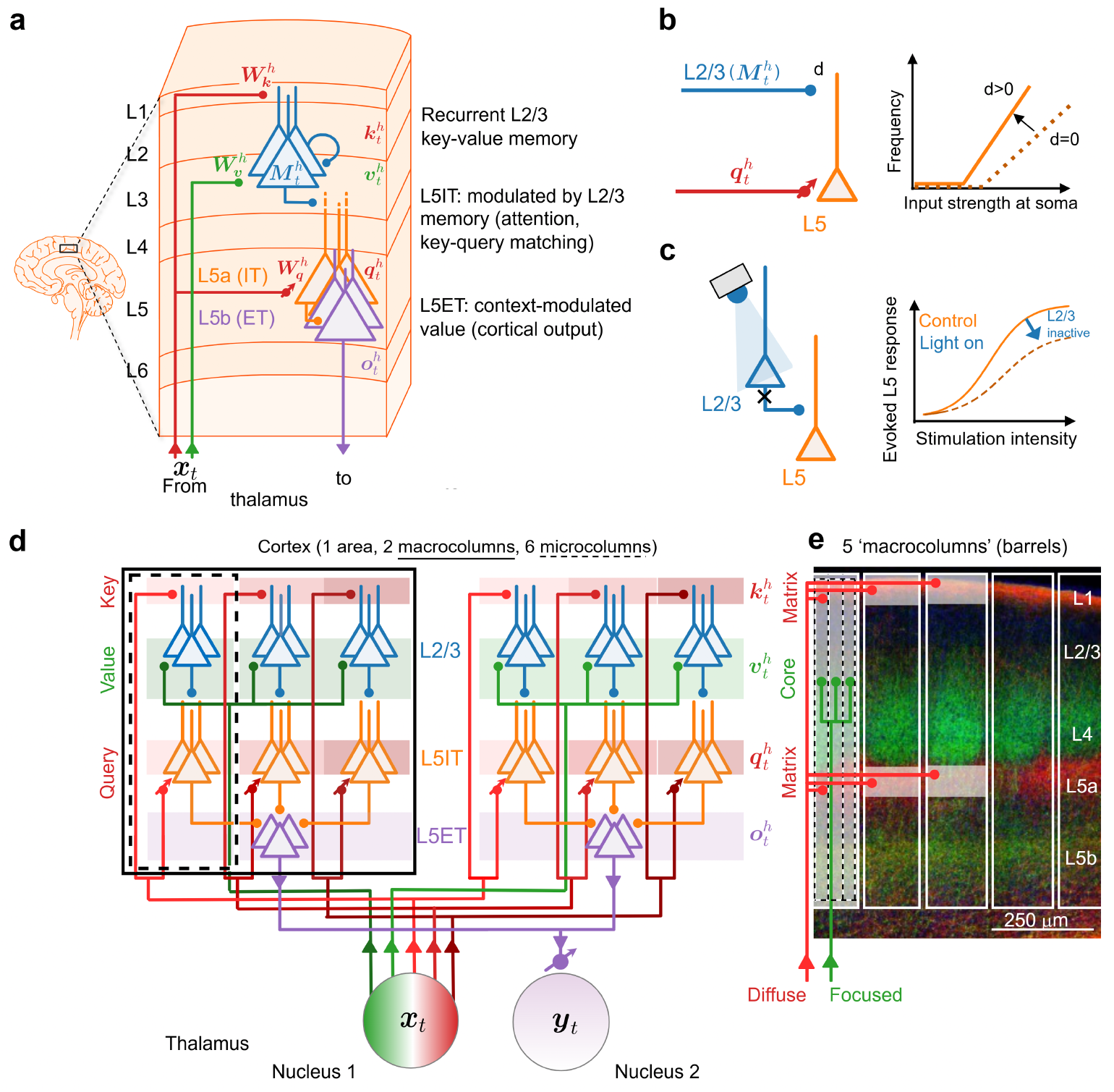
Cortical micro- and macro-columns of layer 2/3 (L2/3) and layer 5 (L5) pyramidal ensembles implement the computation of self-attention. (**a**) A L2/3 pyramidal ensemble (blue) performs component-wise integration of the key–value pairs (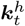, via L1, red; 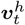, via L3, green) over time (eq. 4). A L5a intratelencephalic (L5IT) pyramidal ensemble (orange) spatially integrates the key-value memories (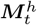 eq. 1 from the L2/3) in their apical dendrites; this apical input modulates the query representation (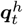, red) arriving at the basal dendrites of L5IT pyramidal neurons. A L5b extratelencephalic (L5ET) pyramidal ensemble (purple) integrates in space the L5IT activity, and sends the cortical output 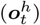 to the thalamus (eqs. 5 and 6). (**b**) Apical input gain-modulates the somatic firing rate of L5 pyramidal neurons elicited by basal (perisomatic) input (adapted from [45]); *d >* 0 and *d* = 0 correspond to the cases with and without apical inputs, respectively. (**c**) The L2/3 pyramidal neurons gain-modulate the activity of L5 pyramidal neurons: photo-inhibition of L2/3 pyramidal neurons decreases the gain of the L5 activity (adapted from [48]). (**d**) Core and matrix structure of thalamo-cortical projections that compute keys, queries, and values. One cortical area (head *h*) with 2 *×* 3 = 6 cortical microcolumns (one shown in panel a), arranged in 2 macrocolumns (*d*_*v*_ = 2, *d*_*k*_ = 3). A thalamic nucleus encodes the current input token ***x***_*t*_ (green-red bottom circle; color shades indicating the thalamic sources of the core- and matrix-projections) and sends core-type afferents focused to L2/3 (green; a line represents a single value component *v*_*i*_), and sends matrix-type afferents diffusively to L1 and L5a (differently shaded red; a line represents a single component of the key *k*_*j*_ and query *q*_*j*_, respectively). The thalamo-cortical projections to the L5IT pyramidal neurons (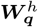, encoding the queries), and the readout connections to the next thalamic nucleus 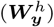 are considered to be plastic (symbolized by the small diagonal arrow crossing the synaptic terminal). (**e**) Fluorescence imaging data for core (green) and matrix (red) thalamo-cortical projections reproduced from Staiger and Petersen [12]. Overlaid is a single afferent fiber of the diffuse matrix projection (red, single component of the key and query vector), distributed horizontally across macrocolumns (5 solid white rectangle), and a single afferent fiber of the focused core projection (green, single component of the value vector) projecting to all microcolumns (3 dashed white rectangles) within a single macrocolumn.

L2/3 pyramidal neurons receive distinct inputs via basal and apical dendrites [44]: we exploit this distinction by assigning value signals 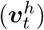 to basal dendrites via synapses 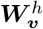, and key signals 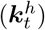 to apical dendrites via synapses 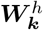 (fig. 1 a). The somatic output is the product of basal and apical activity, 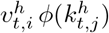, consistent with the observed apical gain modulation of perisomatic activity [45]. Combined with short-range intra-ensemble recurrent connections [39, 40, 41] that integrate somatic activity over time, this yields the recurrent key-value memory,

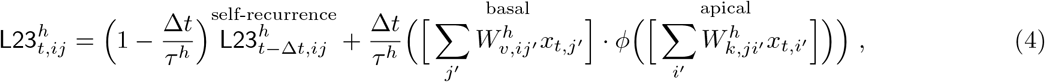

where we replace 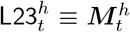(eq. 1) in our neuronal mapping. In eq. 4, 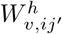 *′* and 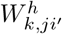 are the basal 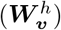 and apical 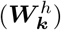 synapses on the postsynaptic L2/3 neuron *ij*, relaying the individual thalamic afferents *x*_*t,j*_*′* and *x*_*t,i*_*′* to the basal and apical dendrites, respectively. The function *ϕ* represents a dendritic nonlinearity.

### Query-based memory retrieval in cortical layer 5 pyramidal neurons

L5 contains two major types of neurons, intratelencephalic (IT) and extratelencephalic (ET) neurons with distinct targets and computational roles [46]. The local pathway from L2/3 to L5 pyramidal neurons is established as a major component of the canonical cortical circuit [47]. Experimental evidence shows that L2/3 pyramidal neurons modulate the gain of L5 neurons [48] (fig. 1 c). We exploit this canonical pathway to implement query-based memory retrieval: L2/3 ensembles project to ensembles of L5IT neurons within the same vertical microcolumn, supplying the key-value matrix 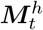 via apical inputs (fig. 1a).

L5 pyramidal neurons, similar to those in L2/3, receive distinct inputs via basal and apical dendrites [49]. We assign query inputs to the basal dendrites of L5IT neurons. There are therefore *d*_*v*_ × *d*_*k*_ L5IT ensembles, with *d*_*q*_ query components distributed *d*_*v*_ times along the cortical sheet. Each L5IT pyramidal ensemble computes the product of its basal query input and its apical memory input,

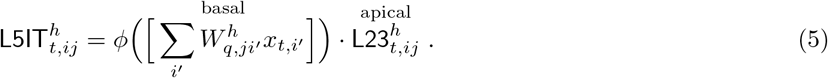

In other words, the activity in the L5IT neurons is the product of L2/3 memory matrix with the diagonal matrix associated with the query, i.e. 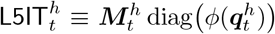. This product can be interpreted as L5IT gain modulation by apical input and by L2/3 neurons, and this gain modulation is itself well documented ([45, 48], fig. 1 b,c).

According to the implementation of the self-attention mechanism in eq. 5, the L5IT pyramidal neuron *ij* is gain modulated by the single L2/3 neuron *ij*. While this matches the formal self-attention calculation, the biology provides multiple apical inputs that ensure robustness against noise. To formalize this, we extend the apical inputs to include adjacent L2/3 pyramidal neurons, 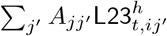. The apical afferent matrix ***A*** is concentrated around the main diagonal and selects the L2/3 pyramidal neurons that project to a specific apical dendrite of a L5IT neuron. Computationally, this leads to the attention signal formed by the scalar product with respect to matrix ***A***, 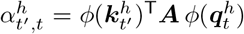. As we show below (fig. 2d), this biological feature makes the key-query comparison more robust against synaptic fluctuations.

**Figure 2:**
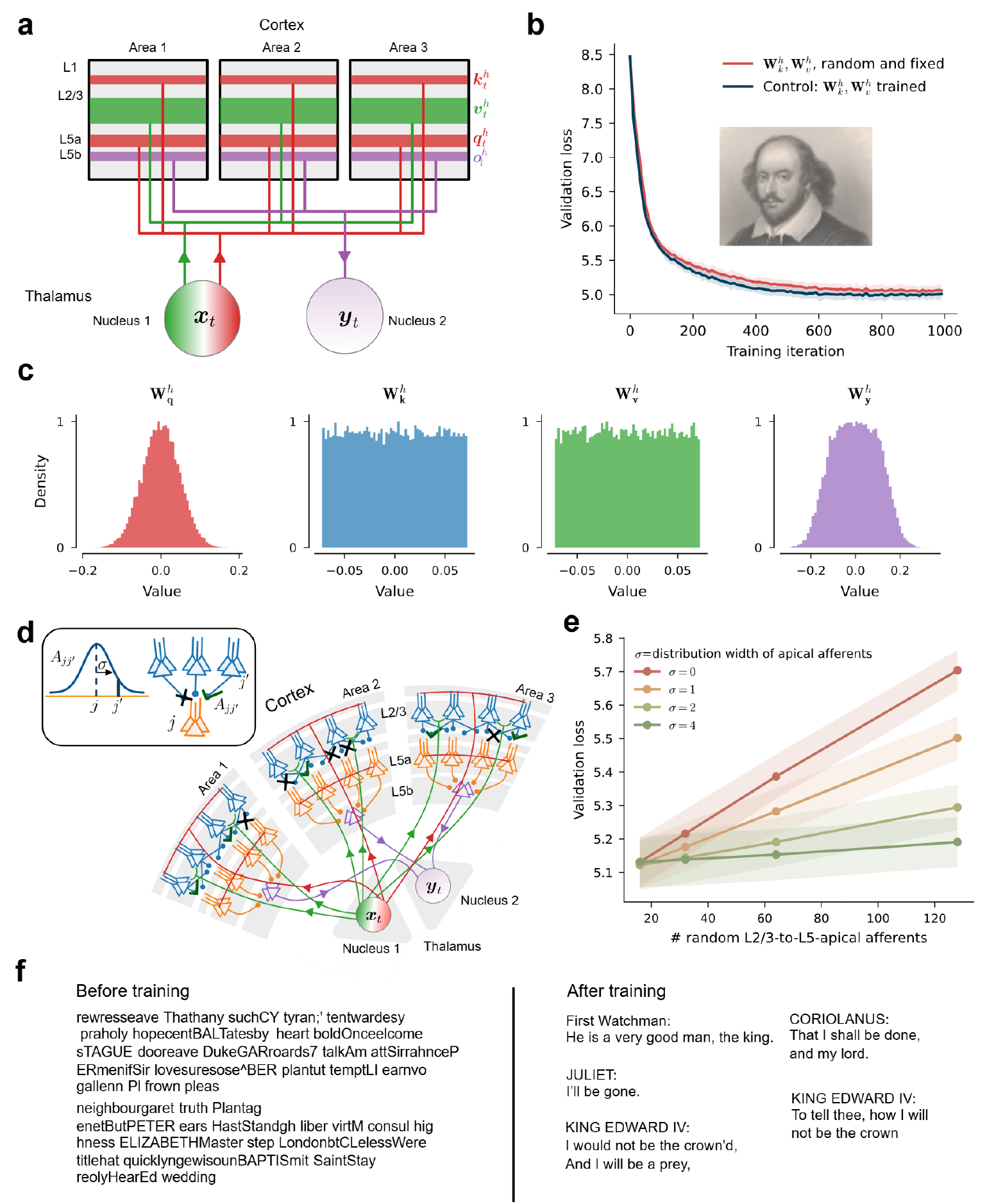
Parallel processing and plasticity in cortical areas generating Shakespeare language. (**a**) Matrix thalamo-cortical projections compute keys (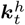, red line in layer 1) and queries (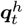, red line in layer 5) out of the input ***x***_*t*_ from nucleus 1. Core thalamo-cortical projections compute values (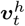, green, L2/3, see also fig. 1). Cortico-thalamic projections from L5ET pyramidal neurons convey the context-modulated values from the different areas, 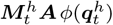, to a higher a thalamic nucleus that computes ***y***_*t*_. (**b**) Validation loss of the cortical MHSA network learned on the Tiny Shakespeare data set, with 6 parallel areas (‘heads’), and each area with 32^2^ = 1024 L2/3 and L5IT pyramidal neurons (red). If in addition the memory synapses 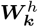 and 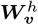 are learned as a control, the similar validation loss is achieved (blue). The mean (solid lines) and standard deviation (shaded regions) are calculated across 10-fold cross-validation. (**c**) Histograms of the learned query and readout synapses (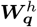 and 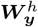) and fixed memory synapses across the 6 cortical areas. Before learning, 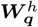 and 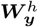 histograms are similar to the memory synapses. (**d**) Cortical circuit showing the randomized dropping (small black crosses) /adding (small green ticks) of L2/3-to-L5IT-apical afferents *A*_*jj*_*′* . (**e**) In the presence of randomized connections (i.e., random ***A*** entries), the validation loss is stabilized with a broader distribution of the L2/3-to-L5IT-apical afferent strength *A*_*jj*_*′* around the diagonal (red: *σ* = 0 with ***A*** the identity matrix; yellow: *σ* = 1, green: *σ* = 4, see text). The mean (solid lines) and standard deviation (shaded regions) are calculated across 10-fold cross-validation. (**f**) Illustrative outputs from the network before (left) and after (right) training.

L5ET neurons then sum the L5IT activity to produce the head output [46]. As the cortical output, L5ET projects further to thalamic nuclei [50]. Hence,

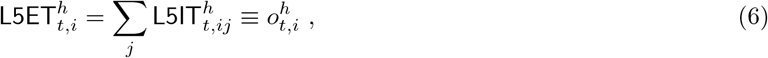

where for clarity of the neuronal mapping we introduced the notation 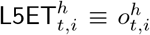 for the output of neuron *i* in head (area) *h* at time *t*. The L5 connectivity implements the matrix multiplication of eq. 2, where the matrix 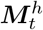 is encoded in the L2/3 activity itself. The outputs from different cortical areas are integrated in higher thalamic nuclei, which project back to the cortex; there, the new tokens are processed in a subsequent cortical self-attention block, forming thalamo-cortico-thalamic chains [51]. In summary, L2/3 pyramidal neurons store key-value pairs over time, while L5 pyramidal neurons represent the context-dependent mixture of values retrieved by the current query (eq. 2).

### Core and matrix thalamo-cortical projections compute keys, queries, and values

The representation of the key-value memory, together with the query-based retrieval, requires specific thalamo-cortical projection patterns that distribute the thalamic input across cortical layers. Intriguingly, these spatial projection patterns have been experimentally observed. The thalamus is a major driver of cortical activity, conveying sensory or higher-order thalamic information through two distinct pathways, the core and the matrix projections [52, 12, 51]. Core projections are focused in space and target L3, directly or via layer 4 [47]. Matrix projections are diffuse in space and target layer 1 and layer 5. These two projection types offer the substrate to represent the outer product between values and keys, 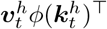, in the cortical area (indexed by the head *h*).

The outer product 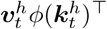 combines each component of the value vector 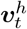 with each component of the key vector 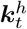, yielding the *d*_*v*_ × *d*_*k*_ grid of L2/3 pyramidal neuron ensembles. The L2/3 neurons are driven by the basal and perisomatic core projections that are focused in space, while being modulated by apical matrix projections that are distributed in space (fig. 1 d,e). A single component 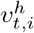 projects to all the *d*_*k*_ microcolumns within a macrocolumn (3 green terminals in L2/3, fig. 1 e, ending in 3 microcolumns of the first macrocolumn). In the somatosensory cortex of the mouse, a macrocolumn corresponds to a ‘barrel’, with neurons in L2/3 and L4 being driven by the input from a single whisker [12]. A single component 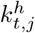 of the key, instead, is distributed to the homologous (the ‘same’) microcolumns within all the *d*_*v*_ macrocolumns of a given area *h* (3 red terminals in L1, fig. 1 e, ending in the first microcolumn of 3 macrocolumns).

Query components 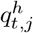 are distributed through diffuse matrix projections to L5a, analogous to the key components in L1. In L5a the matrix projections terminate on the basal and perisomatic compartments of L5IT pyramidal neurons (lower 3 red terminals in fig. 1 e). The L2/3 pyramidal neurons project to the apical dendrites of L5IT pyramidal neurons and modulate their gain (eq. 5). As L2/3 pyramidal neurons integrate the key-value memory across time *t*′ ≤ *t*, an individual L5ET ensemble, 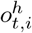, eventually represents an output of one of the *d*_*v*_ macrocolumns of area *h* (eqs. 2 and 6). The first-order thalamic nuclei encode the input tokens, ***x***_*t*_, while the higher-order thalamic nuclei integrate the cortical outputs 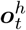 from all heads into the final output ***y***_*t*_ (eq. 3).

### Parallel processing in cortical areas by multi-head self-attention

The cortico-thalamic feedback circuit forms a shallow hierarchy in which multiple cortical areas process, in parallel, different aspects of the same thalamic input([53], fig. 2a). For example, in the visual system, distinct areas simultaneously extract shape, color, and motion from the same scene [54]. The multi-head self-attention (MHSA, eqs. 1 to 3; [4]) offers a framework to incorporate such parallel processing by multiple cortical areas (multiple heads *h*). In our interpretation, a thalamic nucleus projects to several cortical areas in parallel, and each area independently computes its own keys 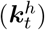, queries 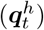, and values 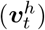 from the same current sequence element (***x***_*t*_, a ‘token’). In natural language processing, such a sequence element ***x***_*t*_ encodes a word or a syllable, and in visual processing it encodes the foveal input from a saccade, for instance. The cortical outputs (eq. 6) are integrated in a higher-order thalamic nucleus [55], computing ***y***_*t*_ that can serve as a prediction of the next sequence element, ***y***_*t*_ ≈ ***x***_*t*+Δ*t*_, if it is a higher-order sensory nucleus, or of a motor output if it is a motor nucleus ([56], see fig. 2a and fig. 3d).

**Figure 3:**
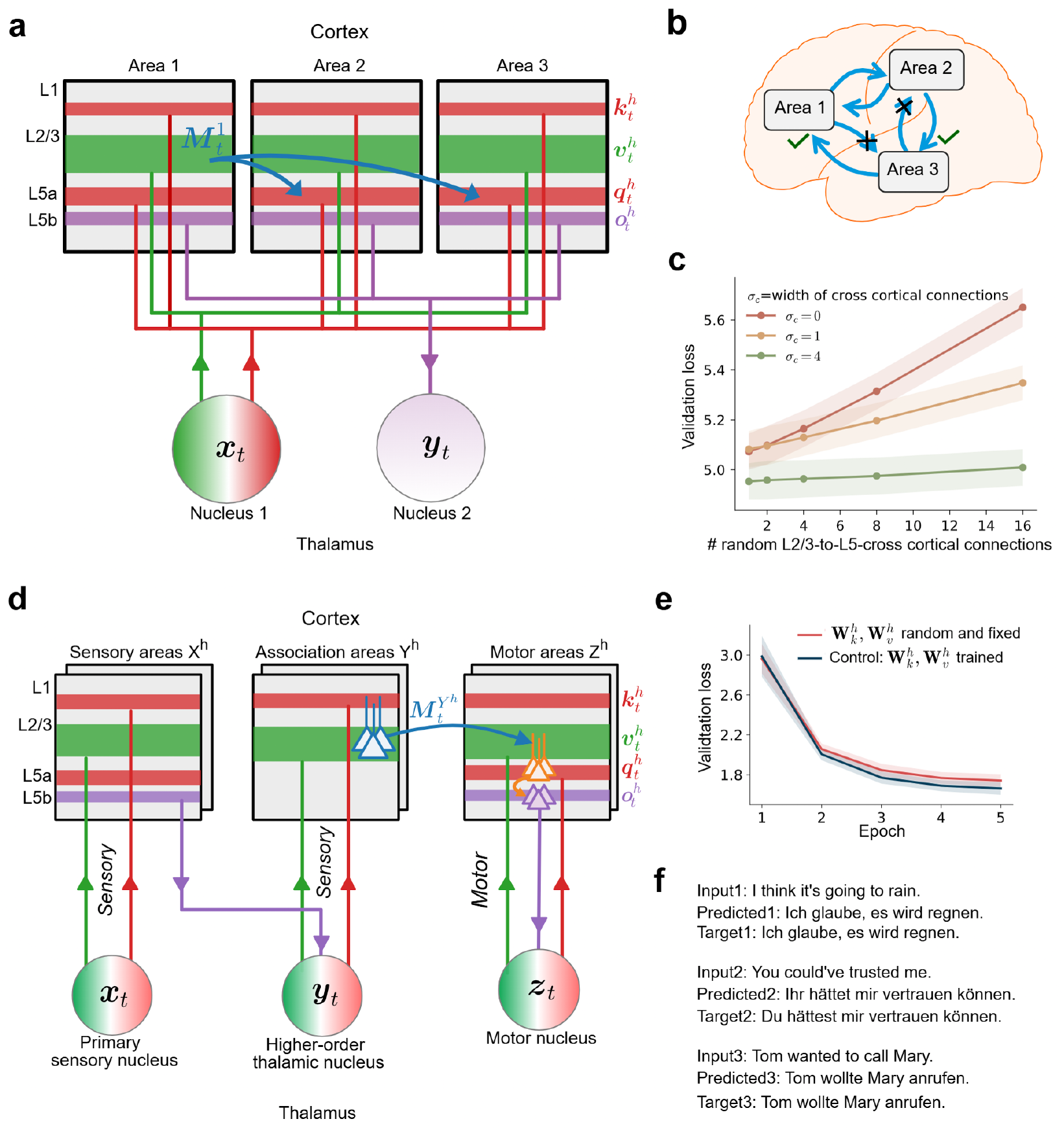
Cross-attention across cortical areas: (**a**) Each thalamic nucleus encodes the current element (***x***_*t*_) and projects to distinct cortical areas (fig. 1 and fig. 2a, and eq. 17). Blue arrows denote cross-connections between different areas (heads) encoding the same sequence (***x***_*t*_). (**b**) A more biological depiction showing randomized connections (dropout: black crosses; dropin: green ticks) between cortical areas. (**c**) Additional cross-connections (eq. 17) between areas provide robustness against the randomized connections. Increasing *σ*_*c*_ increases the strength of the cross-connections between different cortical areas (see eq. 17). The mean (solid lines) and standard deviation (shaded regions) are calculated across 10-fold cross-validation. (**d**) Thalamus-driven multimodal cross-attention: two thalamic nuclei encode the current elements of two different sequences ***x***_*t*_ and ***z***_*t*_, here sensory and motor sequences, and project to sensory and motor cortical areas *X*^*h*^ and *Z*^*h*^, respectively. Projections from nucleus ***x***_*t*_ target the upper cortical layers (L2/3 to L1) and update the key-value memory in sensory areas *X*^*h*^. Higher-order sensory areas (association areas) *Y* ^*h*^ receive driving input from lower sensory areas *X*^*h*^ via higher-order thalamic nucleus ***y***_*t*_ (beside possible modulating cortico-cortical cross-connections as in panel a). Projections from the motor nucleus ***z***_*t*_ target cortical layer 5 (right, the ‘decoder’ pathway in the transformer language [4]) and query the L2/3 sensory memory 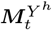 (eq. 8) from the sensory areas (left, the ‘encoder’ pathway; here shown for the higher-order association areas *Y* ^*h*^), in addition to L2/3 motor memory 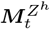 in motor areas *Z*^*h*^. For purely thalamic cross-attention see fig. S1. (**e**) Validation loss of the cortical MHSA network learned to translate English sentences (represented by the ‘encoder’) into German (represented by the ‘decoder’). Both encoder blocks (*X*^*h*^ and *Y* ^*h*^), and also the decoder block (*Z*^*h*^), consist of 6 parallel areas (heads), with each area containing 100^2^ = 10 000 L2/3 and L5IT pyramidal neurons (red). As a control we also learned the memory synapses 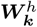 and 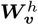 and achieved a similar validation loss (blue). The mean (solid lines) and standard deviation (shaded regions) are calculated across 10-fold cross-validation. (**f** ) Illustrative sentences translated by the multi-head cross-attention network (subsumed in the MHSA abbreviation; Input: English; Predicted and Target: German).

We trained a cortical MHSA network with six heads and a single hierarchical level (a ‘block’) on the Tiny Shakespeare dataset to perform next-word prediction (see Methods and section S7 for pseudocodes). While training, we keep the strength of the key and value synapses, 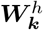 and 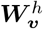, random and fixed, while only learning the query and readout synapses 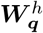 and 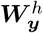. Randomness makes the memory representations for different input elements ***x***_*t*_*′* (*t*′ ≤ *t*) uncorrelated, resulting in a high memory capacity [57], and fixed synaptic strengths remove the need to train the network by error-backpropagation through time (BPTT, or involve additional eligibility traces, see S5). In fact, learning only the query and cortical readout synapses from instantaneous errors does not degrade performance in the generation of Shakespeare language (fig. 2b,c,f, see also [58, 59]). The linear self-attention mechanism we consider here is extensively tested and benchmarked in [36], with all weights (including 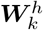 and 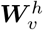) trained.

In fig. 2d, we show the biological substrate of our MHSA network with the retrieval from the L2/3 memory 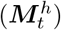 via L5IT and L5ET pyramidal neurons, leading to the readout activity 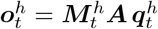. Here we included the L2/3-to-L5IT apical afferent matrix ***A*** introduced above (below eq. 5). Consistent with the topographic organization of cortical columns, we consider this projection matrix to be Gaussian distributed along the diagonal, 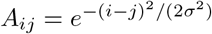, with *σ* controlling the width of the distribution (see inset in fig. 2d). This biological feature makes the validation loss robust against random synaptic perturbations, such as drop-in or drop-out (fig. 2e, Methods), mimicking slow fluctuations in the synaptic strengths that may underlie representational drift in cortex [60].

### Cortico-cortical and thalamo-cortical cross-attention

The cortex is characterized by extensive inter-areal connections [10]. In our framework, cortico-cortical con-nections involve L2/3 pyramidal neurons encoding key-value memory (eqs. 1 and 4) in one cortical area and projecting to the apical dendrites of L5IT neurons (eq. 5) in one or more other cortical areas [61] (fig. 3(a)). Similarly, L5IT pyramidal neurons in a given cortical area can receive apical inputs from L2/3 pyramidal neurons in many cortical areas.

We model cortico-cortical connections via a cross-cortical apical afferent matrix ***C*** (see eq. 17); *C*_*hh*_*′* defines the strength of L2/3 to L5IT apical afferents from cortical area (head) *h* to *h*′. Consistent with the decaying number of cortico-cortical connections with interareal distance [62], we define 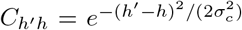. Here, *σ*_*c*_ controls the width of the L2/3-to-L5IT cross-cortical projection pattern, and for monkey cortex *σ*_*c*_ is on the order of a centimeter ([62], with area indexing *h, h*′ = 1, 2, ..). We observe that networks trained with larger *σ*_*c*_ show an increased robustness of the validation loss in the presence of randomized drop-in and drop-out L2/3-to-L5IT cross-cortical projections (fig. 3(b,c) see Methods). The multiple cortico-cortical cross connections go beyond the single cross-attention architecture of transformers [4], and equip cortex with a redundancy code that makes the sensory representation more stable against noise. By summing up the cortical activities (eq. 3), the redundancy further allows for amplifying important sensory features, as has been postulated based on experimental data [63].

The cortico-cortical connections described above include mutual connections between two cortical areas via L2/3-to-L5IT cross-cortical projections. These cortical areas can encode the same or different sensory modalities (fig. 3a, [64]), or they encode sensory and motor information (fig. 3d, [65]), each with multiple heads. To formalize the sensory-motor interactions, we first consider sensory areas *X*^*h*^ which process sensory input tokens ***x***_*t*_ encoded in a sensory thalamic nucleus. The higher sensory (association) areas *Y* ^*h*^ then receive and process the outputs ***y***_*t*_ from the areas *X*^*h*^ (fig. 3d). The motor areas *Z*^*h*^ process motor tokens ***z***_*t*_, which are encoded in a motor thalamic nucleus. The L2/3 memory in a higher sensory area *Y* ^*h*^ is queried by L5IT pyramidal neurons in the corresponding motor area 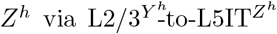 cross-projections from *Y* ^*h*^ to *Z*^*h*^ (see eq. 2 and fig. 3d), producing the output

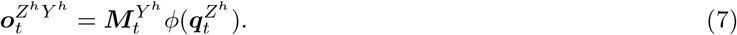

The above output is represented by L5ET pyramidal neurons in the motor area Z^*h*^ (eq. 6) that project to the motor thalamic nucleus where the next motor token 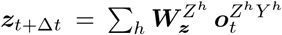 is generated through cortico-thalamic synapses 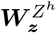 (see also eq. 3). At this next time step *t*+Δ*t*, the motor output is fed back to the motor areas *Z*^*h*^, while the new sensory input ***x***_*t*+Δ*t*_ feeds to the sensory areas *X*^*h*^. The outputs ***y***_*t*+Δ*t*_ from *X*^*h*^ feed into the sensory areas *Y* ^*h*^ to form the memories 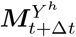, which cross-project to the corresponding motor areas to calculate the next output 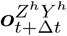.

According to the cross-attention expressed in eq. 7, the sensory L2/3 memory modulates the motor query that is represented in the basal dendrites of L5IT motor pyramidal neurons. But the sensory modulation may also be complemented by the local memory within the motor area itself, with both types of sensory and motor memories (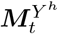 and 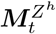) summed up in the apical dendrites of the L5IT motor neurons (fig. 3d). The self- and cross-modulated motor output is then again integrated in the motor nucleus to form the next motor token,

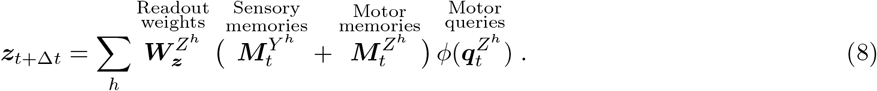

Unlike the standard transformer, which alternate self- and cross-attention layers while forming concatenations of head vectors [4], the suggested cortical self- and cross-attention combines both, while summing the outputs of the cortical motor areas in the thalamic nucleus. Our cross-attention architecture implements an encoder-decoder structure [4], with sensory areas analogous to the encoder and motor areas analogous to the decoder (fig. 3d).

We trained a cortical MHSA (that includes cross-attention) network (Methods) to translate English to German (fig. 3e,f). In our translation task, the English sentence plays the role of the sensory input and the German translation that of the motor output. The auditory thalamus projects English tokens ***x***_*t*_ into auditory cortical areas *X*^*h*^. Sequential word-by-word translation is incorrect in general. For instance, translating “Do you bake bread?” into “Backen Sie Brot?” requires the second-last word, ‘bake’, to begin the translation with ‘Backen’. We therefore presented first the full English sentence from time 0 to *T* in order to encode it in the auditory memories (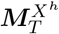 and 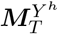). The memories 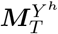 project to motor areas *Z*^*h*^, which also receive the translated German token ***z***_*t*_ from the previous time step; together they produce the prediction ***z***_*t*+Δ*t*_ of the next German token (see eq. 8 and Methods). During training, this prediction was compared with the next German target token, and the query and readout synapses (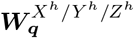 and 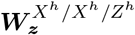) were adapted based on the prediction error (see below), while the key and value synapses were again kept fixed and random. During testing, the next German token becomes the prediction ***z***_*t*+Δ*t*_.

Anatomically, the multi-model cross-attention formulated in eq. 7 can be implemented in multiple ways. We described cross-attention from a sensory cortical area *Y* to another motor cortical area *Z* (fig. 3a), but the two areas may also be the same, forming a single cortical association area receiving thalamic inputs from different modalities *Y* and *Z* (fig. S1).

### Matching to human intracortical activity during speech perception

After training our cortical MHSA network on the translation task, we asked whether cortical activities in patients can be mapped to the model activities while presenting the same English sentences. We compared the auditory embeddings (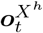 from above) with intracranial recordings obtained from 4 epilepsy patients during perception of short spoken sentences (in a seizure-free period, fig. 4a). For each word heard, the MHSA network provides one embedding per head, and each recording contact provides the broadband high-frequency activity (BHA; 70–150 Hz) at a specific cortical location, a well-validated proxy for local neuronal firing. We mapped a contact onto a head as follows: we reduced each word embedding of each head to a binary label, namely whether it projected more strongly onto the first or the second principal component of that head’s embeddings, and trained a classifier to predict this label from the BHA of the contact (three-fold cross-validation, Methods). If the classification accuracy of a contact–head pair was significantly above chance, the contact was mapped onto that head; a contact mapped with several significant heads was assigned to the head providing the highest classification accuracy.

**Figure 4:**
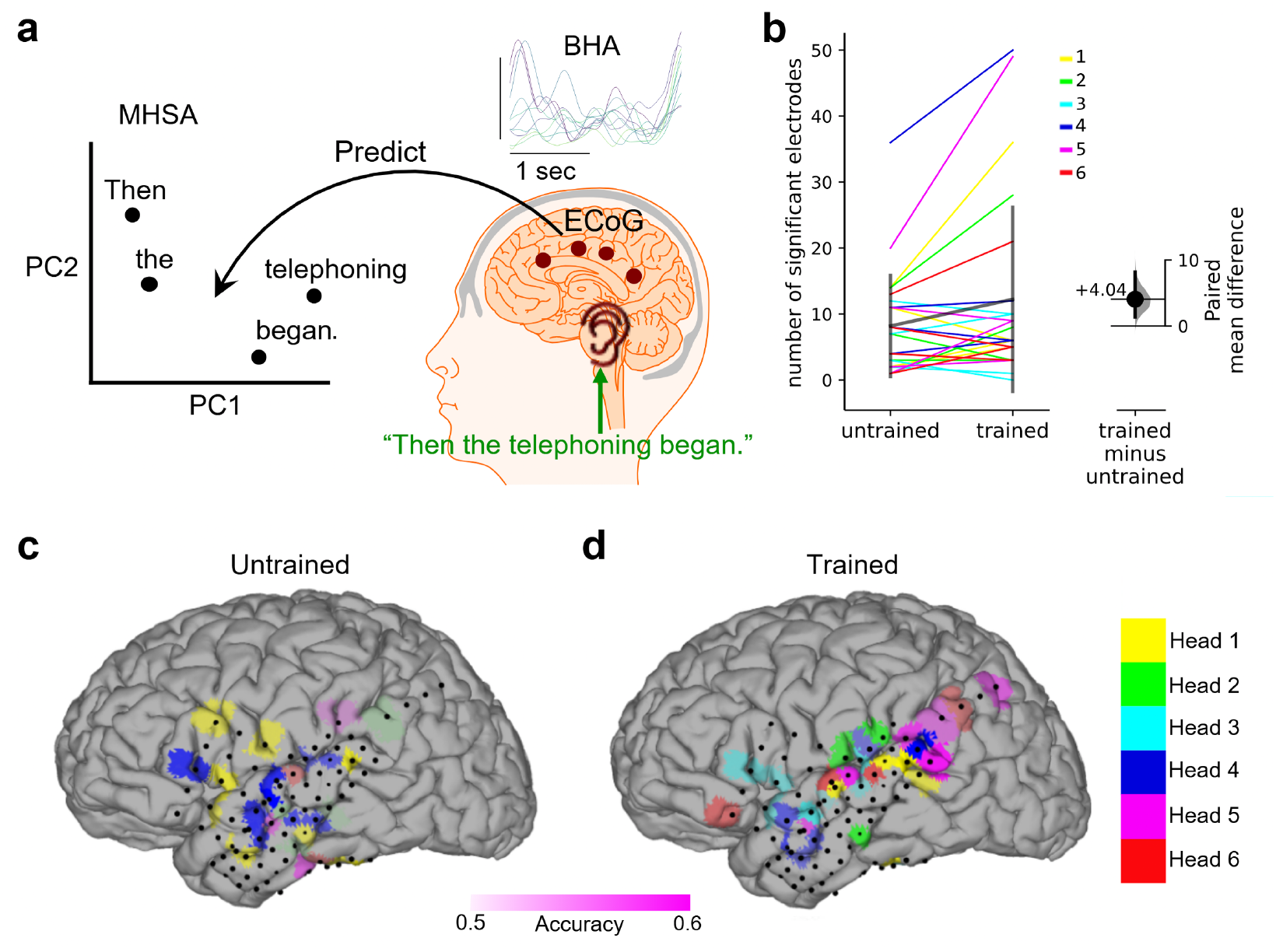
Mapping MHSA networks to human cortical areas with intracranial recordings. (**a**) English sentences (example in green) are presented to the cortical MHSA network and to participants implanted with electrocorticography (ECoG) grids. From the cortical signals (quantified as the broadband high-frequency activity, BHA, a proxy for local neuronal firing rates) we predicted for each word whether the cortical output vector 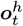 was closer to the first or second principal component (PC_1_ or PC_2_) extracted from all word outputs (Methods). (**b**) Number of recording contacts for which the BHA significantly predicts the PC’s of the MHSA embeddings, for both the untrained and trained models, across the 4 participants. Left: each line corresponds to a single ECoG contact, and each color to a different MHSA head. Right: difference of the number of significantly predicted heads for the trained minus untrained MHSA model. (**c**,**d**) Colors indicate the most predictive head for each significant electrode for the untrained and trained cortical MHSA network for participant 3, with color placed at the cortical region corresponding to the projected nearest-contact position [66]. Only for the trained MHSA network was the positional mapping consistent across validation folds (fig. S2b). The head-area mapping after training (d) reflected the cortical areas known to be involved in speech perception [67]. The accuracy of the prediction is shown as color transparency, scaled between 0.5 (chance level) and 0.6.

For the trained network, more contacts mapped onto a head than for the untrained network, in all four participants (fig. 4b; significance against 1000 label-shuffled surrogates, with correction for false discovery rate, see Methods). Two observations support the resulting mapping. First, it was reproducible: for the trained model, the classification accuracies of a contact across heads were correlated between the three cross-validation folds, i.e. the same heads were consistently predicted with highest accuracy in every fold, whereas for an untrained model, the accuracies varied from fold to fold (fig. S2b). Second, it was anatomically coherent: the mapped contacts clustered along the superior temporal gyrus, in line with classical models of speech perception [68, 67]. Contacts in different cortical areas were assigned to different heads, so that each area was preferentially mapped to one head (figs 4d and S2a). For the untrained model, in contrast, the mapped contacts were sparser and spatially scattered across brain regions (fig. 4c). Overall, the data confirm that different cortical areas can be consistently mapped to different heads of a MHSA network during a cognitive task.

### L5ET gain-modulation by the unsigned reward-prediction error (RPE)

Our cortical model, so far, learned to predict the next input token. In biology, however, predicting the input is often only a subordinate goal towards the more relevant goal of predicting reward or punishment in the future. We propose a further selection of cortical outputs at the level of L5ET pyramidal neurons and their elaborate apical dendrites ([71], eqs. 2 and 6) via unsigned reward-prediction error (RPE) calculated in subcortical structures, as it has been experimentally observed [23].

Reward prediction and reinforcement learning in the brain is mainly attributed to the basal ganglia and the midbrain [16, 72]. Cortico-basal ganglia feedback loops are involved, for instance, in reward-based adaptation of movements [73, 74], while neurons in the motor cortex and the associated thalamic nuclei carry prospective signals about upcoming reward-directed actions [75]. The basal ganglia receive cortico-striatal inputs from L5IT neurons [76], and via midbrain and thalamus sends a reward-related salience signal back to cortex [77]. In our model, basal ganglia estimate the discounted future reward and a RPE, identified as temporal difference error 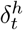 (Methods, [78]). Our basal ganglia-cortical feedback projects to the apical dendrites of L5ET neurons in order to enhance reward-relevant cortical output (figs 5a and 6a). Dopaminergic neurons in the midbrain are well known to encode prediction errors, and a significant fraction is shown to encode a salience signal as unsigned 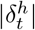 [79]).

**Figure 5:**
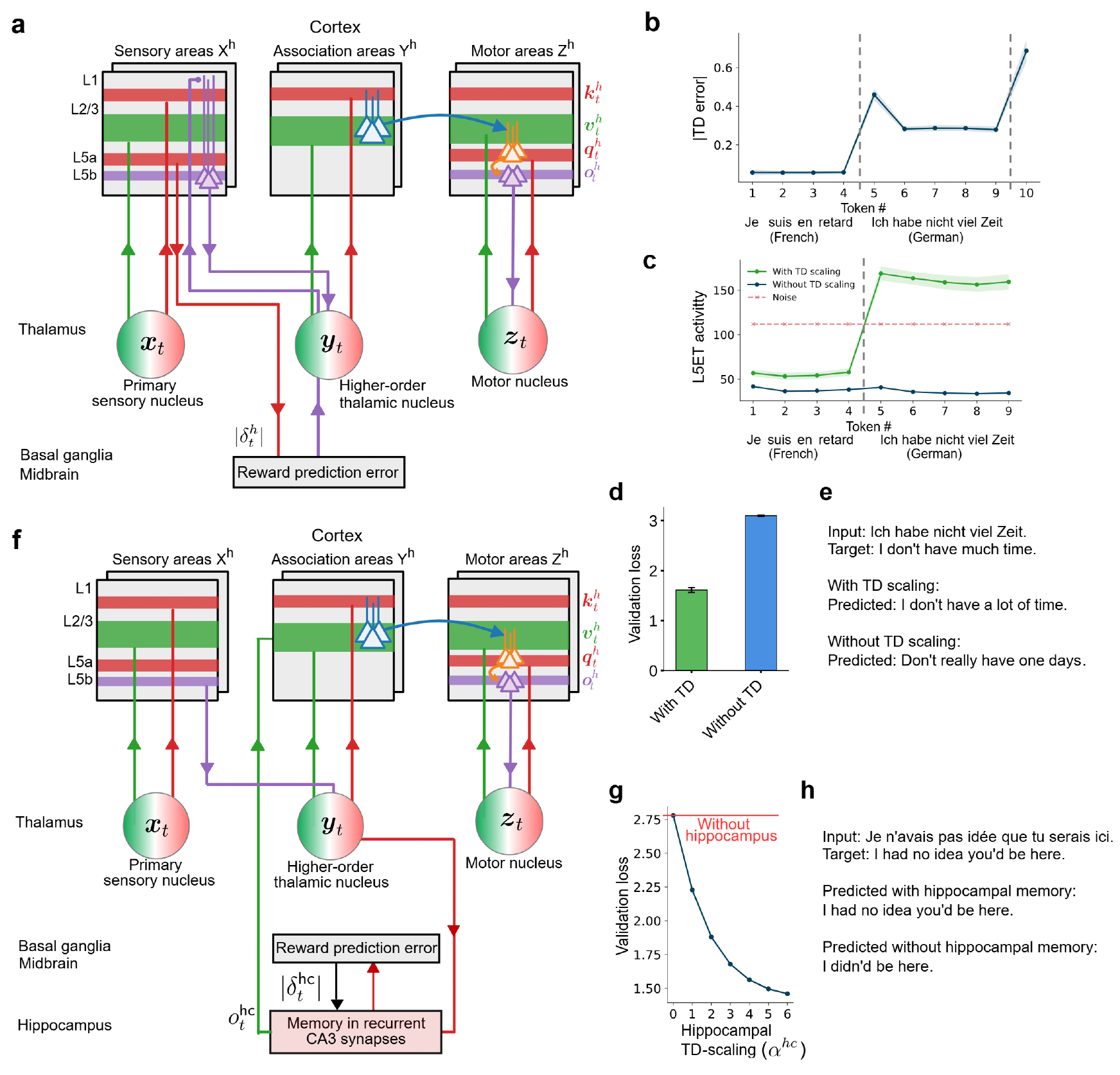
Reward-based modulation of cortical L2/3 memories and L5b output by basal ganglia and hippocampal memories during language translation. (**a**) The cortico-basal ganglia feedback loop for reward-prediction error. L5IT pyramidal neurons in sensory areas *X*^*h*^ project to the basal ganglia, which in turn gain-modulate L5ET pyramidal neurons in areas *X*^*h*^ by the unsigned RPE (the absolute value of the temporal difference (TD) error, 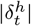, eq. 20, via L1). For a close-up see fig. 6a. (**b**) The mean of unsigned RPEs across heads 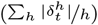 on validation data, for each token of the French and German sentence aligned to the token-axis (mean and standard deviations of 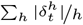 across 10 training runs). During training, the German-English translation was rewarded by its quality, but no reward was given after French sentences. (**c**) Mean absolute component value of L5ET activity eq. 6 (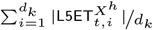 averaged across heads and across tokens in validation data set with the same position within a sentence) without noise term (only first term in eq. 9, solid lines) in the presence of the TD scaling (*α*=6, green) and without TD scaling (*α*=0, blue). With TD-gain modulation, the L5ET signal is above the noise level (mean absolute component value of noise in eq. 9, red dashed line), without TD-gain modulation, it is below. (**d**) A lower validation loss is achieved for German sentences with TD-gain modulation (green) as compared to without (blue) in the presence of noise. (**e**) Illustrative sentences for German-English translation with and without TD-gain modulation. (**f** ) Higher-order thalamic nuclei (encoding ***y***_*t*_) recall hippocampal memories stored in CA3 synapses [69, 37], and project back to higher cortical areas (*Y* ^*h*^, [70]), modulated by the unsigned RPE conditioned on the hippocampal memory. The synaptically encoded hippocampal memory is reactivated in the L2/3 memory of the higher cortical areas (green upwards arrows). (**g**) Increasing the coupling strength of the hippocampal TD error (*α*^*hc*^ in eq. 11) decreases the validation loss for the French-English translation. Horizontal red line shows the validation loss in the absence of hippocampal memory. (**h**) Illustrative sentences for French-English translation, with temporal difference (TD) scaling for gating the hippocampal memory input.

The unsigned RPE is postulated to sharpen the sensory representations in areas *X*^*h*^ via distal apical inputs to L5ET pyramidal neurons [77, 23, 80], see fig. 5a and fig. 6a. With the unsigned 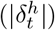 calculated in the basal ganglia-midbrain loop, the activity of the L5ET pyramidal neurons in the sensory areas (eqs. 2 and 6) becomes gain-modulated according to

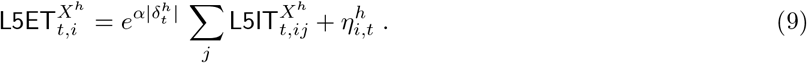

**Figure 6:**
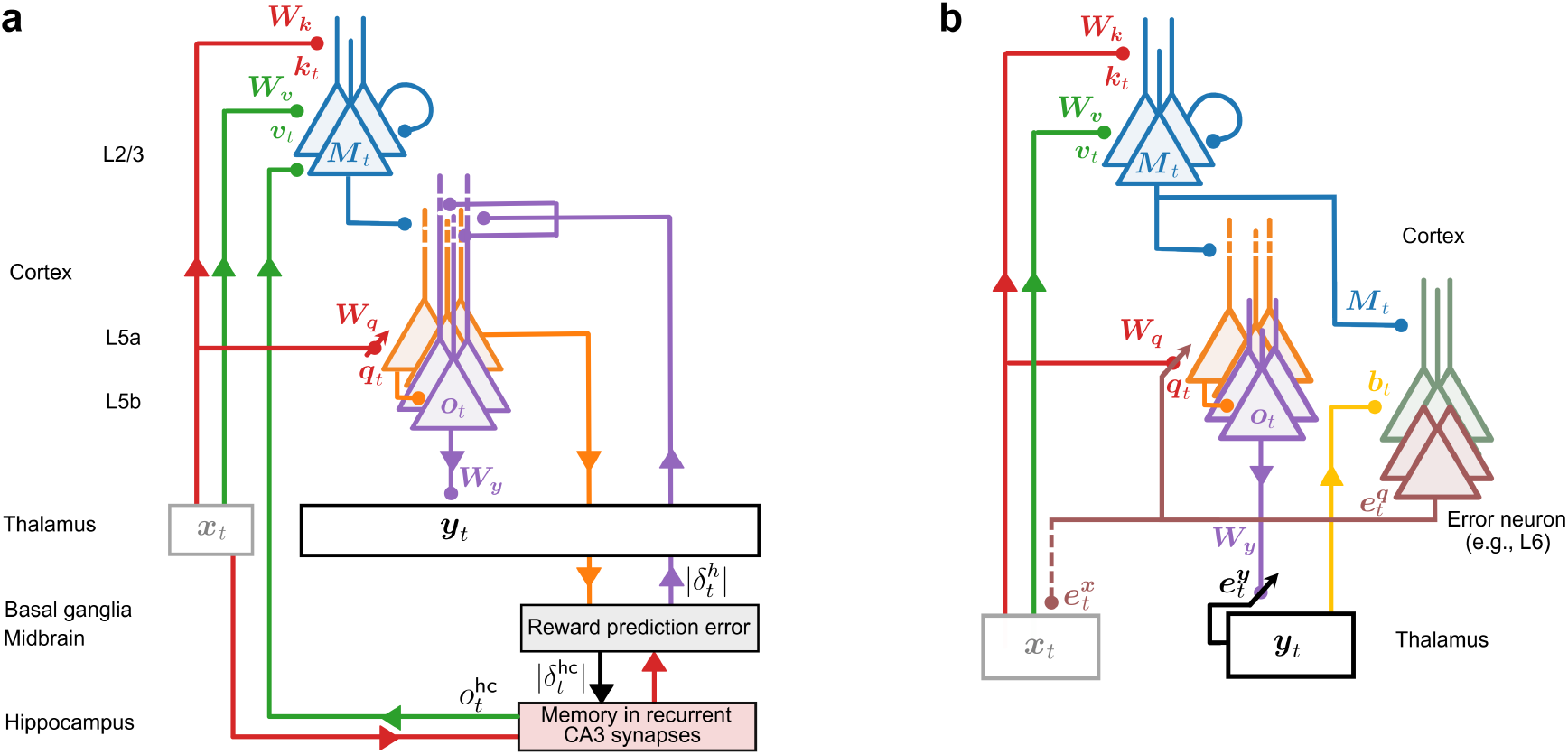
Cortical microcircuits for inference and plasticity. (**a**) L5IT pyramidal neurons (orange) project to basal ganglia and midbrain in order to estimate future reward, and the unsigned RPE is fed back to the apical tuft of L5ET pyramidal neurons to modulate their gain (purple). Hippocampal memory (red transparent) is recruited from the thalamus, in our simplification (red, matrix-type), and feeds back to L2/3 neurons, in our model to selectively (gated by the RPE) extend the neuronal memory by hippocampal synaptic memory. (**b**) Error backpropagation from thalamus to cortex and back. The thalamic prediction error 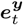 (black) directly modulates plasticity of the cortico-thalamic synapses (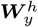, eq. 12; *h*-superscript dropped in the figure). The thalamic error is backpropagated to (e.g., [83]) L6 error neurons as 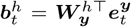 (yellow) where it forms the memory-modulated error 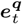 that gates the error-correcting plasticity of the query synapses (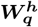, eqs. 13 to 15). The query error 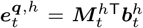 represented in the L6ET pyramidal neurons is further backpropagated to the preceding thalamic nucleus to form the sensory prediction error 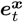(brown dashed, see eq. S35) and gate the gradient-based plasticity in this previous kernel (putatively projecting further back to cortex).

The exponentiation can be seen as apical nonlinearity, and the *α* as basal ganglia-cortical coupling strength. We also added the Gaussian noise 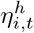 to the L5ET response. The salience modulation is postulated to be specific to each area (index *h*), yet the same for each macro-column within that area (no index *i*). The learning of the discounted future reward is based on minimizing the somato-dendritic prediction error for neurons nudged, say in basal ganglia, by the reward signal ([81], Methods).

To illustrate the computational benefit of the salience-based sensory amplification, we make the language translation task more challenging. First, we present our RPE-modulated MHSA network streams of French and German sentences, and require for the German sentences only to be online translated, while ignoring French sentences. Upon a German-English translation, we provide a reward signal to the basal ganglia that captures the quality of the translation ([82], Methods), while not giving any feedback upon French sentences. Second, we add noise to our sensory representation at the level of L5ET pyramidal neurons, as expressed in eq. 9.

The German-English translation is only possible if the basal ganglia feedback is strong enough to pull the cortical representation of the German out of the noise level (fig. 5b,c; eq. 9). After learning the readout, query and the basal-ganglia synapses, the unsigned 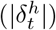 signals the presence of the German words, with a stronger signal at the onset and offset of the German sentence (fig. 5b). This reflects the fact that, when German is presented, more reward or punishment can be harvested as compared to the average across all inputs that include non-rewarded French sentences. The simulations confirm that the unsigned RPE can be seen as a motivational or salience signal that sharpens the sensory representation at the level of the cortical output by gain-modulating L5ET pyramidal neurons [23, 80]. In the absence of this salience signal, there is no reasonable translation, and the validation loss remains high (fig. 5d,e).

### Selecting cortico-hippocampal memories by unsigned RPEs

Our L2/3 key-value memories are stored in the neuronal activities, not in the synaptic strengths as is typically considered (see e.g. [37]). To still recall old memories, they are retrieved from the synaptic long-term stores and made available in the neuronal working memory.

Such a two-stage memory retrieval has two advantages. First, it allows for a selective memory re-activation from the long-term store, made available for motor actions. Second, and more importantly, the memories must also modulate gradient-based plasticity in other synapses, and this is only possible if the memories are available in neuronal activities that can inform plasticity in these other synapses (see below).

As an example of re-activating synaptic memories into L2/3 neuronal memories, we focus on recalling hippocampal memories to higher sensory areas *Y* ^*h*^. The reinstantiation of hippocampal synaptic memories as cortical neuronal memories makes specific hippocampal contents available for read-out from cortex, connecting our proposal to hippocampal index and consolidation theories [17, 18]. Instead of calculating the value 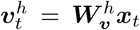 for the cortical area *h* out of the input token ***x***_*t*_, the value may be delivered from the hippocampal (Hopfield-type) memory [37]. For this, the hippocampal memory is itself queried with 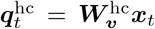, so that it returns the output

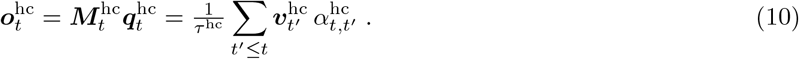

as input to the basal dendrites of the cortical L2/3 pyramidal neurons (Methods, eq. S3). Because the hippocampal memory 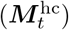 is stored in recurrent CA3 synapses [37] and persists much longer than the L2/3 working memory (encoded in the neuronal activity), we set the hippocampal decay factor to *λ*_hc_ = 1 for simplicity.

The hippocampal input 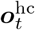 is added to the value vector 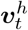 generated from the thalamic input in the L2/3 basal dendrites, and it is attached to the cortical key, 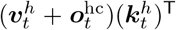 (suppressing the dendritic nonlinearity *ϕ* for convenience). When querying the L2/3 memory at time *t*, not only old values 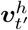 are retrieved (eq. 2), but also values 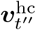 stored in hippocampus at time *t*^*′′*^ ≪ *t*′ further back. This is because the L2/3 memory ***M***_*t*_ stores past key-value pairs, 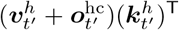, while the hippocampal output 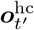 consists of a sum of even older attention-weighted value vectors 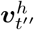 for *t*^*′′*^ = *t*′ −Δ*t, t*′ −2Δ*t*, … (eq. 10, evaluated at *t*′). Overall, generalizing the representation in eq. 2, the readout of the combined cortico-hippocampal working memory at the level of a L5ET pyramidal neuron in the cross-attended area *Z*^*h*^ becomes

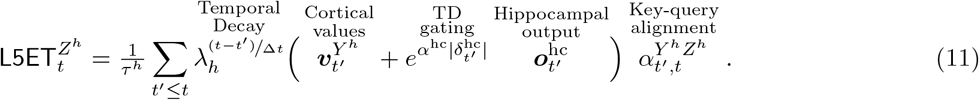

We introduced an additional weighting of the hippocampal memory by the unsigned 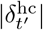, so that the memory is included only when it promises more reward or punishment than expected (Methods, eq. 21). In eq. 11, we omitted the self-attention term within area *Z*^*h*^ for simplicity; see eq. 22 for the full expression for the translation task.

As opposed to adding memories in parallel, as in eq. 8, the hippocampal and cortical memories in eq. 11 are stacked serially in time (see also figs 5f and 6a). Stacking the value-key-query principle from neurons to synapses, and potentially to the encoding of gene transcriptions for the induction of synaptic plasticity (see section S3), can considerably extend the memory time horizon. Re-writing the hippocampal memory into the active L2/3 memory, weighted by the unsigned RPE (eq. 11), reduces the validation loss in a German-English translation task (fig. 5g). The stronger the contribution of the longer-term hippocampal memory (higher *α*^hc^), the better the translation performance (fig. 5g,h).

### Error-correcting plasticity via layer 2/3 memory

Key-value memory is represented in neuronal activities rather than synaptic strengths. The computational justification for these neuronal activities is that they are directly involved in the gradient-descent plasticity rule. To implement gradient-descent learning on squared-error cost function (eq. S12) in neuronal terms, we start with the thalamic error 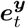 representing the error of the network output. The error represents the deviation of the sensory prediction ***y***_*t*_ from a target, 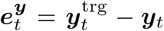, with target given by the next input token, 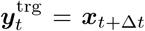. Such thalamic prediction errors are found in behavioral experiments, jointly with sensory prediction error in the cortex [19]. The thalamic error drives the error-correcting plasticity of the cortico-thalamic readout synapses according to (fig. 6b)

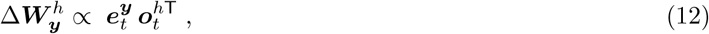

telling that the change of the readout weights 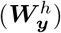 is the product of the (somato-dendritic) prediction error 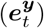 times the presynaptic activity 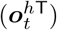 [33].

The thalamic prediction error is backpropagated to the sensory cortex through the (transposed) readout weights and the memory matrix, 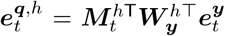, forming the error for the query representation ***q***^*h*^ in area *h* (for convenience we again consider a linear dendritic transfer function *ϕ*). Correspondingly, the gradient-descent plasticity of query synapses is (fig. 6b, section S5)

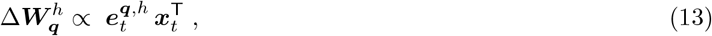

which is of the form of a local error times the presynaptic activity.

The error of the query representation 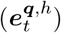 can be further ‘backpropagated’ to the preceding thalamic nucleus (eq. S34) where it adapts the readout of the corresponding earlier sensory areas. That preceding thalamic error can itself be backpropagated again into the earlier sensory areas, forming a chain that propagates errors and instantaneously adapts readout and query synapses throughout. The two gradient rules (eqs. 12 and 13) confirm that no BPTT is required to train our MHSA network for which the synapses of the key-value memory are randomized and frozen.

### Biological implementation of the gradient plasticity

The plasticity of the cortico-thalamic readout weights 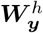 (eq. 12) can readily be implemented in terms of a somatic nudging by the thalamic error signal 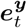, and plasticity being driven by the somato-dendritic prediction error [33, 34]. We next explain how the brain may implement the query error 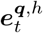 entering the plasticity rule for the query synapses 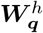 (eq. 13). The crucial insight is that this requires cortical error-backpropagation of the type introduced in [31, 32], 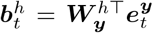, with a gain-modulation of the backpropagated error 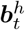 in the analogous way as our L5IT pyramidal neurons are gain-modulated by the L2/3 input (figs 1c and 6b). In fact, the query error is 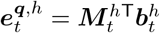, taking the analogous form of our cortical output 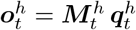.

In analogy to the L5 signal representation (eqs. 5 and 6), we propose an explicit cortical representation of errors, e.g., in L6 pyramidal neurons, which have been shown to be sensitive to surprise [83]. We consider L6 error neurons represented by intra-telencephalic (L6IT) pyramidal neurons. These L6IT pyramidal neurons integrate in their basal dendrites the backpropagated thalamic errors 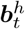 (fig. 6b, yellow), while being gain modulated by the L2/3 pyramidal neurons representing the components of the neuronal memory matrix ***M***_*t*_,

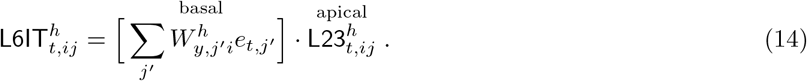

The L6 extra-telencephalic (L6ET) error neurons integrate the activity of L6IT error neurons, 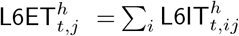 . In matrix form we have 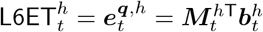.

On the one hand, these L6ET error neurons project peri-somatically to L5IT pyramidal neurons in order to nudge them with the query error 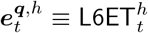 and gate plasticity of the query synapses 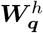 on their basal dendrites (fig. 6b, brown, see also eq. 13),

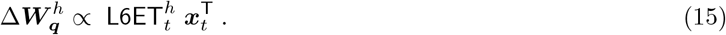

On the other hand, the L6ET error neurons project back to the preceding thalamic kernel to provide the sensory representation ***x***_*t*_ a sensory prediction error 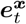 (fig. 6b, brown, see also eq. S34). There is a formal duality between inference and learning (section S6) that is also expressed in the analogy between the L5 output calculation and the L6 error calculation.

The L6ET projection pattern is corroborated by anatomical studies showing that some axonal L6ET projections end in L5a [13], while others end in the preceding thalamic nucleus [51, 84]. Such cooperative thalamo-cortical circuits for sensory prediction errors are also observed during sensory-motor learning [19]. Consistent with experimental findings, we further assumed that instantaneous errors are represented in the thalamic nuclei [85, 86] and modulate synaptic plasticity of our thalamo-cortical L5IT 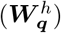 and cortico-thalamic 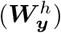 synapses [87, 88, **19]**.

## Discussion

Motivated by the simplicity of the classical key-data associative memories [35] and their revival as high-performance self-attention networks [4, 36], we mapped MHSA networks to the anatomy of thalamo-cortical projections and the physiology of cortical pyramidal neurons. In our framework, L2/3 pyramidal neurons implement an active associative key-value memory that is queried by L5 pyramidal neurons. The gain modulation of L5 neurons by L2/3 memory neurons [48] is the crucial operation that results in the context-modulated cortical output. Different cortical areas correspond to the different attention heads of a transformer block, whose outputs are integrated in a higher thalamic nucleus. The widespread cortico-cortical projection pattern generalizes the concept of cross-attention, according to which one cortical area queries the memory in another area. The experimentally observed dendritic modulation of L5 pyramidal neurons by the unsigned RPE [23] helps to re-activate task-relevant memories across cortex and hippocampus. Applied to sensory-motor streams, our RPE-modulated MHSA network learns online to translate German into English in the presence of perturbing activity fluctuations, while also being exposed to French.

For clarity of the self-attention mapping, our model omits layer 4, the thalamo-cortical entry stage. L4 could readily be incorporated as a multilayer perceptron (MLP, perhaps with skip-connections) between the thalamic input and the cortical self-attention block [4]. The remaining softmax operation is well-studied in neuroscience and neuromorphic engineering [89] and has been described as a canonical computation in terms of soft winner-take-all over dendritic activities [90] and divisive normalization [91, 92].

Our cortico-thalamic MHSA network is well suited for scaling, benchmarking against neural and behavioral datasets, and systematic comparison with larger artificial and biological network models of the brain [93]. The alignment of artificial neural network models with neuronal data saturates as model size increases, even when more data are provided, whereas behavioral alignment continues to increase [94]. In turn, large-scale, biophysically realistic brain simulations align better with neuronal data as model size increases, but their behavioral alignment remains limited [95, 96, 97]. We combined a top-down and bottom-up approach to draw on the best of both: proven computational principles from artificial networks that solve hard cognitive tasks (MHSA), together with features of cortical pyramidal neurons (such as gain modulation) and layered thalamo-cortical circuits that can implement those principles.

We proposed an RPE-modulated MHSA architecture that points beyond the classical transformer of AI [4, 36, 6]. Reward-predicting value functions calculated in the basal ganglia and the midbrain, together with their RPEs [78, 16], enter our model in two forms. First, the unsigned RPE sharpens the signal-to-noise ratio at the cortical output via gain modulation of L5ET neurons. Second, the unsigned RPE selects synaptic memories in other brain regions for re-activation in the L2/3 memory.

The activity in the L2/3 network can be seen as a working memory with a relatively short time horizon (minutes), since it is carried by reverberating neuronal activity [98]. Silent synaptic long-term memories from other brain regions, or other neuronal populations within the same region, may be recruited and written into the L2/3 working memory, depending on whether these memories predict more future reward or punishment than average [99]. We considered the example of synaptic hippocampal memories selected by the unsigned RPE and providing episodic context. But synaptic cortical memories, for instance those encoded in the prefrontal cortex [100], may likewise be selectively recalled into sensory L2/3 working memory and provide more abstract context. Our L2/3 working memories distributed across cortex and linked through cross-attention can be seen as a global neuronal workspace [101]. It can be reconfigured on the fly, drawing in selected association and abstraction levels according to the current sensory input, context, task, and reward function.

Thanks to the key-value memory available in neuronal activities, it becomes possible to implement the gradient-based synaptic plasticity via L6-mirrored L5 circuitry (fig. 6). Our postulated L6IT error neurons are modulated by L2/3 memory neurons and integrated into L6ET error neurons. This configuration mirrors the L5IT and L5ET pyramidal neurons that simultaneously carry out the inference, modulated by L2/3. This duality between learning and inference makes the cortical MHSA network attractive for neuromorphic chip design, where in-memory processing and in-memory learning can be combined on a single chip [102, 103]. Mapping learnable MHSA networks onto cortical and sub-cortical architectures becomes also relevant for clinical applications with the recent development of flexible chips for chronic neural brain-machine interfaces (BMIs, [104, 105]). As we showed, it is enough to learn the readout and query synapses, while the key-value memory synapses can remain fixed, and hence no BPTT is required.

Given the performance of our RPE-modulated MHSA networks with their cellular underpinning, we may ask how diseases of the mind can be understood in the framework of inappropriately expressed self-attention. For instance, epileptic seizures might reflect an overly strong L2/3 recurrence that causes self-exciting dendritic bursting [106]; schizophrenia, an overly strong self-attention to specific past memories at the expense of current sensory input [107]; and symptoms of Parkinson’s disease, an aberrant ‘vigor’ that controls the movement gain [73, 108]. Connectome-constrained brain network models have already demonstrated predictive power for an individual’s recorded brain imaging signals such as EEG, MEG or fMRI [2, 3]. A promising next step is therefore to embed our cortical MHSA networks in such a large-scale brain simulation environment, e.g. The Virtual Brain (TVB, [1]) with a realistic 3D connectome, to calibrate the network on existing data [109], and to study the neuronal dynamics, cognitive abilities and behaviour of the healthy and the diseased brain model [2], thereby bridging brain activity and brain function.

## Methods

### Shakespeare task

For the Shakespeare task (fig. 2 and fig. 3c), we used a dataset containing approximately 1 million characters [110]. We used a Byte Pair Encoding (BPE) tokenizer [111] to extract the most frequent 4096 sub-words of the Shakespeare dataset. We embed this vocabulary of 4096 sub-words (tokens) into the token embedding space ***x***_*t*_ ∈ ℝ^192^. For this, each token is identified by a 4096-dimensional one-hot token vector, e.g. ***κ***_*t*_ = (1, 0, 0, …, 0)^T^ ∈ ℝ^4096^ for the first token, and so on. We consider a linear embedding ***x***_*t*_ = ***Eκ***_*t*_ of these tokens with matrix ***E*** ∈ ℝ^192*×*4096^ that is optimized by end-to-end gradient descent on a cost function defined below (all vectors are considered as column-vectors throughout the text).

The embedded tokens ***x***_*t*_ (mean-centered and normalized) are used as inputs to our cortical MHSA network. In the Shakespeare task, our cortical MHSA network consisted of a single attention block with 6 heads. Given the output ***y***_*t*_ ∈ ℝ^192^ (eq. 3), we obtained the next embedded token ***x***_*t*+Δ*t*_ by sampling in the large token space with probabilities ***p***_*t*_ ∈ ℝ^4096^ and projecting back to the embedding space. The sample probabilities are obtained from the ‘residuals’ ***x***_*t*_ + ***y***_*t*_ (mean-centered and normalized) by applying a linear transformation followed by the softmax, ***p***_*t*_ = softmax ***P*** (***x***_*t*_ + ***y***_*t*_) ℝ^4096^, where ***P*** ℝ^4096*×*192^. The *i*^th^ entry of ***p***_*t*_ gives the probability for choosing the *i*^th^ token ***κ***_*t*_ (with a 1 at position *i*) at time *t*. After having chosen this next token ***κ***_*t*_, the embedded next token is ***x***_*t*+Δ*t*_ = ***Eκ***_*t*_. Both matrices ***E*** and ***P*** were learned end-to-end. We consider a linear dendritic transfer function *ϕ*

For learning the ***E*** and ***P***, and also MHSA network parameters (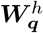 and 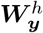), we considered the correct target token 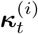 from the training sequence, with index *i* being the *i*’the token in the list of 4096 possible tokens (and hence giving the position of the 1 in the one-hot target vector 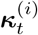. We then calculated the gradient of the cross-entropy cost with respect to the above parameters. The cross-entropy cost at time *t* is obtained by picking out the corresponding component *i* of the sample probabilities,

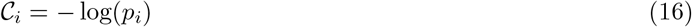

where for convenience we drop the subscript *t*. To optimize the parameters in our network, we averaged the cross-entropy cost over 32 time steps, ⟨*C*_*i*_⟩ = −⟨log(*p*_*i*_) ⟩, where ⟨·⟩ denotes the averaging over the batch of size 32. We trained our network for 5000 epochs using the Adam optimizer [112], and used the early stopping method to reduce overfitting. In fig. 2b we present the training and validation loss for our network.

In fig. 2e, we present the validation loss versus the number of random L2/3-to-L5IT-apical afferents. We defined the (intra-cortical) apical afferent matrix as 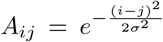 (see after eq. 6) and trained our cortical MHSA network with different values of *σ*. In fig. 2e, we selected the number of random L2/3-to-L5IT-apical afferents (*N* ) as follows: we randomly permute the entries of ***A*** by iteratively exchanging each element, starting from the first, with any other element (including itself) with uniform probability. We then select the first *N* sites and set each of them to 0 or 1 with equal probability.

In fig. 3c, we present the validation loss versus the number of randomized cross-connections between the heads of the same modality via the cross-cortical apical afferent matrix 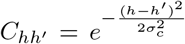 . The matrix ***C*** defines an additional global gating input from the L2/3 memory in area *h*′ to the apical dendrites of L5ET pyramidal neurons in area *h* (beside the gating by the unsigned reward-prediction error from the basal ganglia on L5ET neurons). Formally,

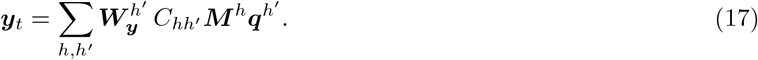

We trained our cortical MHSA network for varying values of *σ*. In fig. 3c, we randomly choose elements of ***C***_*hh*_*′* and set them to 0 or 1 (using the same procedure described for varying ***A***).

### Translation task

For the English to German language translation task (fig. 3e and f), we used approximately 1.6 × 10^5^ sentence pairs (roughly first-half of the English-German sentence pairs in the ManyThings database [113]) in our total dataset. We used a Byte Pair Encoding (BPE) tokenizer to extract the most frequent 16834 sub-words for the English sentences. Each token is identified by a one-hot vector, and we obtain the embedded tokens ***x***_*t*_ for English tokens from their respective one-hot vectors as previously described. English embedded tokens ***x***_*t*_ serve as inputs to sensory areas *X*^*h*^ (details in text). Their outputs are integrated to produce ***y***_*t*_, and the residual ***x***_*t*_ + ***y***_*t*_ (mean-centered and normalized) is passed to areas *Y* ^*h*^. German embedded tokens ***z***_*t*_ are obtained similarly to the English tokens and input to areas *Z*^*h*^, which produce output 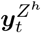. The residual 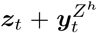 (mean-centered and normalized) is then used to sample the next token ***z***_*t*+Δ*t*_. We consider a threshold-linear dendritic transfer function *ϕ*.

For the plot of the training and validation loss in fig. 3e, we trained an MHSA network with six heads in the encoder and decoder layers. We used a token embedding dimension of *d* = 600 for both the English and German tokens ***x***_*t*_. Keys, queries and values, including the cortical output 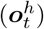, have the dimension *d*_*k*_ = *d*_*q*_ = *d*_*v*_ = 100. We train our MHSA network over 5 to 6 epochs with a batch size of 16, minimizing the cross-entropy loss function (see eq. 16). We split the dataset into training and validation sets in an 8 : 2 ratio.

### Value and TD-estimates via basal ganglia and midbrain

To implement temporal difference (TD) error-based modulation in our MHSA network (fig. 5b and c), we use a similar dataset to the one described above (approximately first 1.2 × 10^5^ English-German sentence pairs in the ManyThings database [113]), but replace a randomly chosen 30% with French sentences. From these sentences we considered the most frequent vocabulary of size 16834 to construct the token embedding as described above. We train our MHSA network to translate German sentences into English and to remain silent when French sentences are presented. We trained our MHSA network over 4 epochs with a batch size of 16. We split the dataset into training and validation sets in an 8 : 2 ratio.

Abbreviating 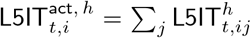 for the L5IT activity in the sensory areas *X*^*h*^, we learn for each head the estimates 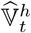 of the value function [114],

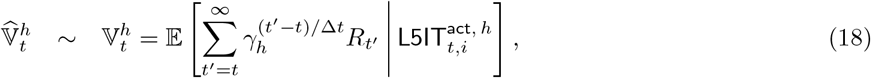

where *R*_*t*_ is the external reward signal, that is vanishing for time steps *t* within a sentence. At the end of a translated sentence, *R*_*t*_ is computed as the character n-gram F-score (chrF) [115] between the translated output and the ground-truth sentence (otherwise *R*_*t*_ = 0) for German to English translation. *R*_*t*_ is always zero for the input French translation.

We model the basal ganglia neurons to learn the estimate 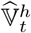 based on the dendritic prediction of somatic activity [81]. With dendritic input 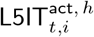 and synaptic weights 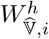, the activities of the value neurons become

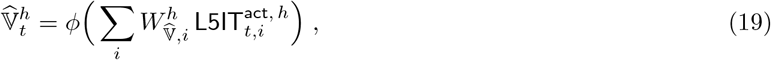

with sigmoidal transfer function *ϕ*. During learning, the soma is nudged by the reward signal *R*_*t*_, such that the somatic voltage becomes 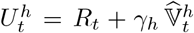, with *γ*_*h*_ being the dendritic attenuation factor (that becomes the discount factor for future reward estimates). The somato-dendritic prediction error is then just the temporal-difference (TD) error [114],

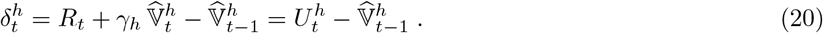

The weights 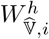 are learned by gradient descent on the squared somato-dendritic prediction error, 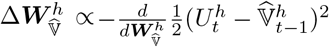. In our framework, we assume that the basal ganglia output to the apical dendrites of the L5ET^*h*^ pyramidal neurons represents the (amplified) unsigned TD-error 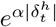.

The noise 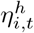 we were adding to the L5ET activities in the translation task (eq. 9, fig. 5c-d), was of the form 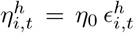, with 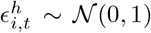 normally distributed, and with scaling factor *η*_0_ = 140. We set *α* = 6 and clipped 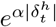 when this expression exceeds 6. We set *γ*_*h*_ = 0.7. In fig. 5c, the mean value of the magnitude of signal components is 161 (comparable to the noise amplitude *η*_0_) after scaling. We first trained the translation task without basal ganglia and noise, then trained the basal ganglia, and during the validation we combined the basal ganglia feedback with the noise.

The TD-values 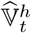 were learned based on a selection of 10% of the sentences from the training dataset described above. We randomly sampled 100 sentences from this set, presented them sequentially, trained 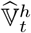 for 100 epochs, and repeated this 1.64 × 10^2^ times for the training data set. We ignored the final punctuation marks during our training. We used the entire validation data set (fig. 5b and c) described above.

### Value-based inclusion of hippocampal memories

To implement TD error-based modulation with hippocampal memories in our MHSA network (fig. 5f,g,h), we trained our MHSA network to translate to English when the French sentences are presented and to repeat when the German sentences are presented. We replace 30% of French sentences (1.2 × 10^5^ sentence pairs; roughly first-half of ManyThings database [113] by German sentences. We trained our MHSA network using the procedure described previously.

For the hippocampal memory 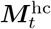 (eq. 10), we learned the estimate

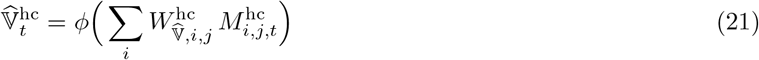

of the value function 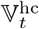 and extracted the hippocampal TD-delta 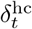 as in eq. 20, but with 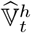 replaced by 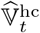.

To prevent instability during training to predict values based on the hippocampal memory input (eq. 21), we used a training procedure with a primary and a target network. The parameters 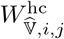 (eq. 21) of the primary network are trained to minimize the temporal-difference error (eq. 20) between the predicted values 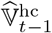 provided by the primary network and the bootstrap targets 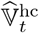 generated by the target network. The target network parameters 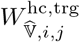 are updated using Polyak averaging (updates following [116]) as 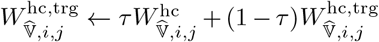 with *τ* = 0.005. The target network provides stable targets during training, as its parameters are updated more slowly than those of the primary network.

For the translation task, the full expression containing the cross- and self-attention with hippocampal memory input (eq. 11) is

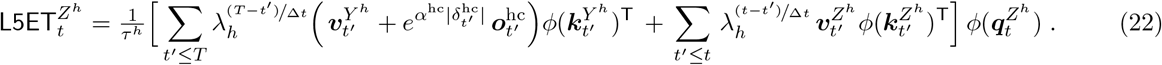

In the above equation, the first term represents the cross-attention, where the memory is formed by the complete input sentence (length *T* ) in area *Y* ^*h*^, and the query is in area *Z*^*h*^. The second term in eq. 22 represents the self-attention within area *Z*^*h*^.

We trained our primary network as follows: from the training dataset (see above), we selected approximately 10% of the sentences. We then randomly sampled 4 sentences from this set and trained for 100 epochs over these sentences sequentially. We repeated this 3 × 10^3^ times for the training data set. We used the entire validation data set (fig. 5g and h) described above. In Fig. 5g, the red line (without the hippocampus) corresponds to the hippocampal output scaled by a factor of 7 × 10^−3^; the network was trained without this factor. This scaled hippocampal output is then amplified by the TD error (eq. 10).

### Participants, auditory stimuli, and data acquisition

The study involved four participants (4 women, mean age 25.6 years, range 19–33) with pharmacologically resistant epilepsy (focal seizures), who underwent subdural electrode array implantation for neuromonitoring of epileptogenic brain areas as part of their clinical epilepsy treatment. The electrode array locations were based solely on the requirements of the clinical evaluation. All participants provided informed consent, and the experiment was approved by the local ethics committee (Albany Medical College Institutional Review Board). No monetary compensation was given to the participants.

Participants passively listened to 112 unique sentences from a transcribed and aligned continuous speech corpus (TIMIT) [117]. Participants were native English speakers with normal sensory and cognitive functions and demonstrated left-hemisphere language dominance.

The implanted ECoG grids (Ad-Tech Medical Corp., Racine, WI; PMT Corporation, Chanhassen, MN) consisted of platinum-iridium electrodes (4 mm in diameter with 2.3 mm exposed) embedded in silicon. The inter-electrode distance was 4 or 10 mm. ECoG signals were recorded using seven 16-channel g.USBamp biosignal acquisition devices (g.tex, Graz, Austria) with a sampling rate of 9600 Hz. The reference and ground were chosen by selecting ECoG contacts that were far from epileptic foci and regions of interest. Data acquisition and synchronization with task stimuli were performed using the BCI2000 software. The participant’s voice was also acquired through a dynamic microphone (Samson R21s) which was rated for voice recordings (bandwidth 80-12000 Hz, sensitivity 2.24 mV/Pa), and placed 10 cm away from the patient’s face. Another dedicated 16-channel g.USBamp amplifier was dedicated to acquiring and digitizing the microphone signal to guarantee synchronization with the ECoG data.

Each patient’s pre-implant high-resolution structural MRI scan was automatically segmented and parcellated using Freesurfer (http://surfer.nmr.mgh.harvard.edu/). Then, a post-implantation high-resolution CT or MRI scan was coregistered with the preimplant MRI scan. Electrode artifacts were identified visually on the post-implantation scan. Electrode coordinates were corrected for the brain shift caused by the implantation procedure by projecting them back to the pre-implant leptomeningeal surface.

### Signal processing, classification task and surrogates

The signals from the ECoG grids were first filtered to remove line noise using a notch filter at 60 Hz and its harmonics (120, 180, and 240 Hz). Common-average referencing was applied. For each channel, we extracted BHA in the 70-150 Hz range [118]. BHA was computed as the average z-scored amplitude of eight bandpass Gaussian filters, with center frequencies and bandwidths increasing logarithmically and semi-logarithmically, respectively. The resulting BHA was down-sampled to 100 Hz. Each word onset was identified and aligned to the neural data. The neural data was then divided into 5 non-overlapping bins of 50 ms each, ranging from 0 to 250 ms after word onset, and averaged within each bin. This resulted in five neural data points per word.

The MHSA cortical output 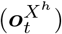 were obtained for each word of the speech stimuli. The MHSA output for each word is obtained by averaging the outputs of its sub-words (tokens), yielding a token per word (of dimension 100). Then, the dimensionality of the MHSA embeddings for each head was reduced to the two leading principal components (PC_1_ and PC_2_). Each word was labeled according to the largest projection of its cortical output 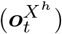 onto the two PC’s, i.e., by the index *i* with the larger scalar product 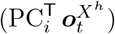 among *i* = 1, 2, and this label was the target of the word classifier. The classifier was trained to predict the target from the 5 neural BHA data points (a number for each of the 5 non-overlapping bins) defined above. The classification was performed independently for each electrode-head pair. We used leave-one-out three-fold cross-validation, the resulting accuracy was averaged across the three folds.

Statistical significance was assessed by comparing the true classification accuracy (ACC) against the classification accuracies obtained for a surrogate distribution. Each surrogate was obtained by reshuffling the labels of the classification task (PC_1_ versus PC_2_). For each surrogate, a surrogate classification accuracy was obtained by performing the same three-fold classification task described above. We thus obtained a null distribution of classification accuracies ACC^∗^ across surrogates. The empirical p-value was then calculated as *p* = (1+#(ACC^∗^ *>* ACC))*/*(1 + *N* ), where *N* = 1000 and #(ACC^∗^ *>* ACC) describes the number of shuffle surrogates which had a higher accuracy than the true classification accuracy ACC. We applied the false discovery rate (FDR) correction for multiple comparisons [119] with chosen targets of *α* = 0.3. The test was performed for each electrode and both the non-trained and trained MHSA embeddings separately. The electrode-head mappings in fig. 4c and fig. S2 are obtained by identifying for each electrode the most significant head with the best classification accuracy. Each head is assigned a color, determining the coloring around the significant electrodes. The triangles of the mesh surface within 10 mm radius of the significant electrode position are colored according to the color of the assigned best head. These colored mesh triangles are projected to the brain surface using the NeuralAct package [66].

## Acknowledgments

We thank Nicolas Deperrois for simulations in the early project stage, and Huifang Wang for comments on the figures. We also thank Robert T. Knight and Peter Brunner for kindly providing the human intracortical recordings.

## Author’s contributions

The research was designed by WS, VB and AG. Mathematical derivations were performed by AG, VB and WS, simulations by VB, and the analysis of the intracortical human data by TP. These 4 authors created the figures and wrote a draft of the manuscript and the Supplementary Materials. MW, TB and AS were engaged in early simulations and discussions, and together with VJ in the final writing.

## Supplementary Materials

**Table 1:**
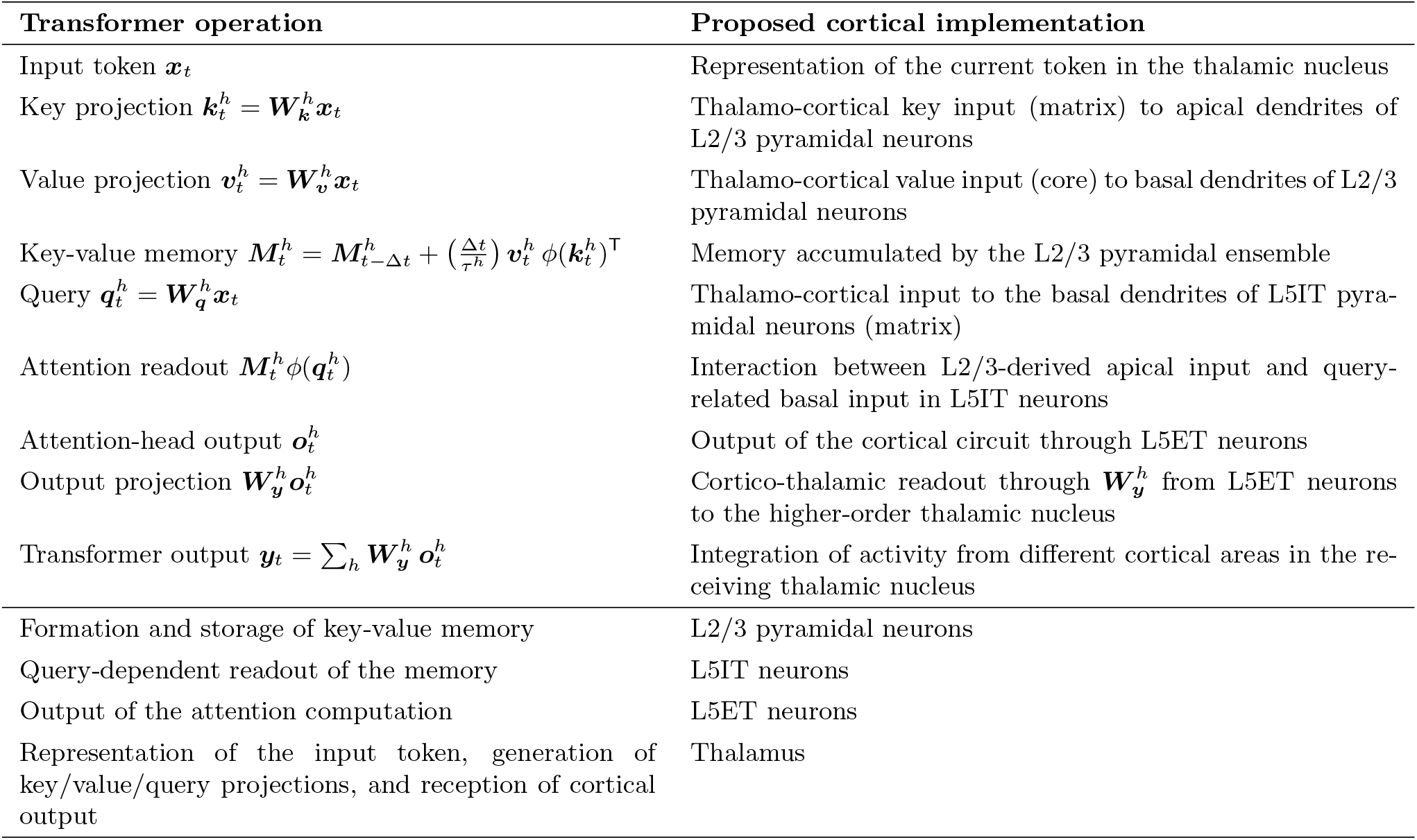
Mapping between transformer operations and proposed cortical implementations.

| Transformer operation | Proposed cortical implementation |
| --- | --- |
| Input token $\mathbf{x}_t$ | Representation of the current token in the thalamic nucleus |
| Key projection $\mathbf{k}_t^h = \mathbf{W}_k^h \mathbf{x}_t$ | Thalamo-cortical key input (matrix) to apical dendrites of L2/3 pyramidal neurons |
| Value projection $\mathbf{v}_t^h = \mathbf{W}_v^h \mathbf{x}_t$ | Thalamo-cortical value input (core) to basal dendrites of L2/3 pyramidal neurons |
| Key-value memory $\mathbf{M}_t^h = \mathbf{M}_{t-\Delta t}^h + \left(\frac{\Delta t}{\tau^h}\right) \mathbf{v}_t^h \phi(\mathbf{k}_t^h)^\top$ | Memory accumulated by the L2/3 pyramidal ensemble |
| Query $\mathbf{q}_t^h = \mathbf{W}_q^h \mathbf{x}_t$ | Thalamo-cortical input to the basal dendrites of L5IT pyramidal neurons (matrix) |
| Attention readout $\mathbf{M}_t^h \phi(\mathbf{q}_t^h)$ | Interaction between L2/3-derived apical input and query-related basal input in L5IT neurons |
| Attention-head output $\mathbf{o}_t^h$ | Output of the cortical circuit through L5ET neurons |
| Output projection $\mathbf{W}_y^h \mathbf{o}_t^h$ | Cortico-thalamic readout through $\mathbf{W}_y^h$ from L5ET neurons to the higher-order thalamic nucleus |
| Transformer output $\mathbf{y}_t = \sum_h \mathbf{W}_y^h \mathbf{o}_t^h$ | Integration of activity from different cortical areas in the receiving thalamic nucleus |
| Formation and storage of key-value memory | L2/3 pyramidal neurons |
| Query-dependent readout of the memory | L5IT neurons |
| Output of the attention computation | L5ET neurons |
| Representation of the input token, generation of key/value/query projections, and reception of cortical output | Thalamus |

### S1 Multimodal thalamo-cortical cross-attention

**Figure S1:**
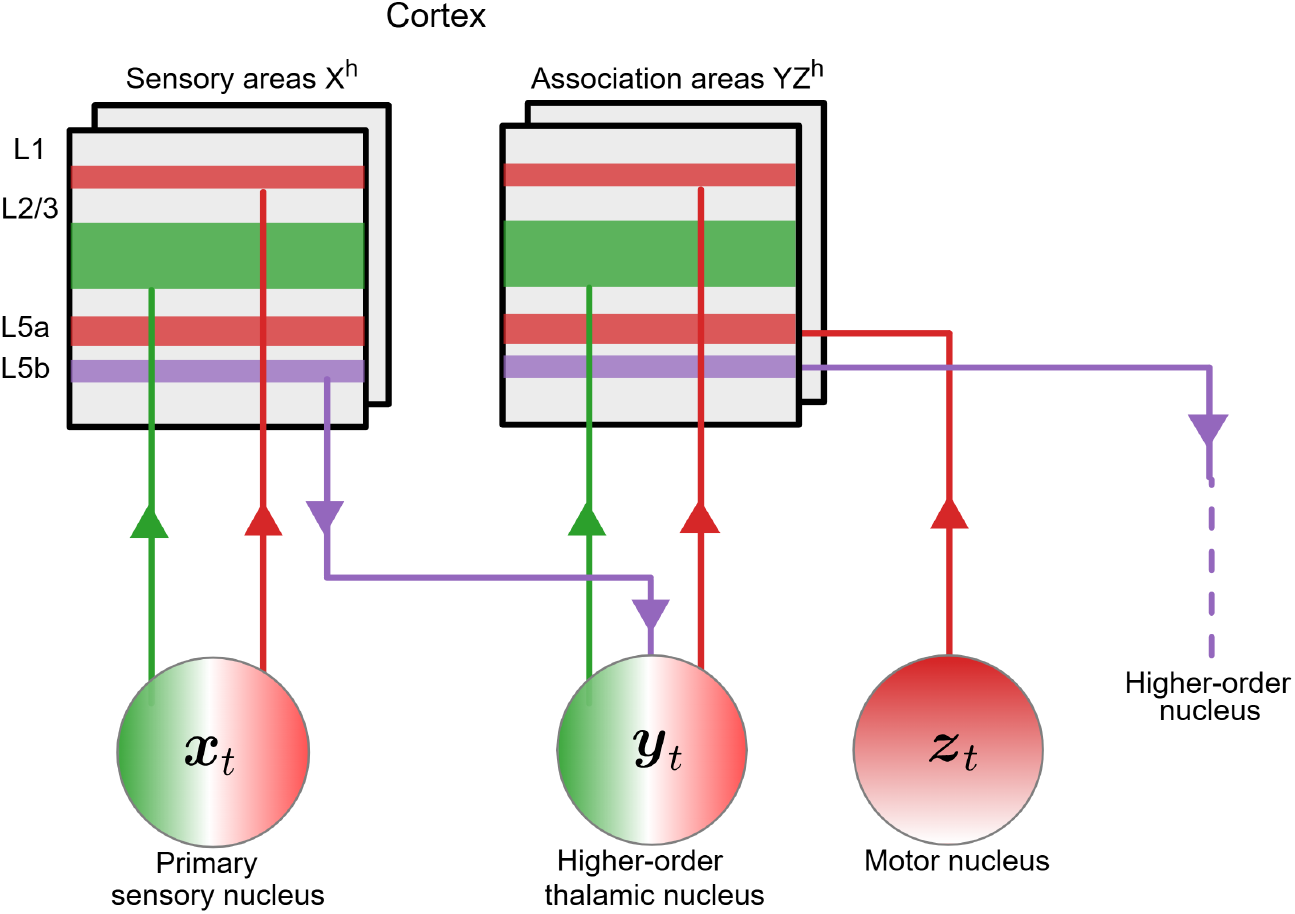
Thalamus-driven multimodal cross-attention in an associative cortical area. The primary sensory nucleus encodes ***x***_*t*_ and projects to primary sensory (auditory) areas *X*^*h*^. The outputs from the areas *X*^*h*^ are integrated in the higher-order auditory thalamus, encoding ***y***_*t*_. Two auditory and visual/motor thalamic nuclei encode the elements of two different sensory modalities, ***y***_*t*_ and ***z***_*t*_, respectively, and project to the same cortical association area [64, 120]. Projections from the auditory thalamic nucleus encoding ***x***_*t*_ target the superficial cortical layers (including L1) of the cortical association area and update the L2/3 key-value memory. Projections from the visual thalamic nucleus encoding ***z***_*t*_ target cortical layer 5a and query the L2/3 memory formed by input ***y***_*t*_ from the other modality. The output from the L5ET pyramidal neurons in L5b projects to a next higher-order (multi-sensory) thalamic nucleus [56].

**Figure S2:**
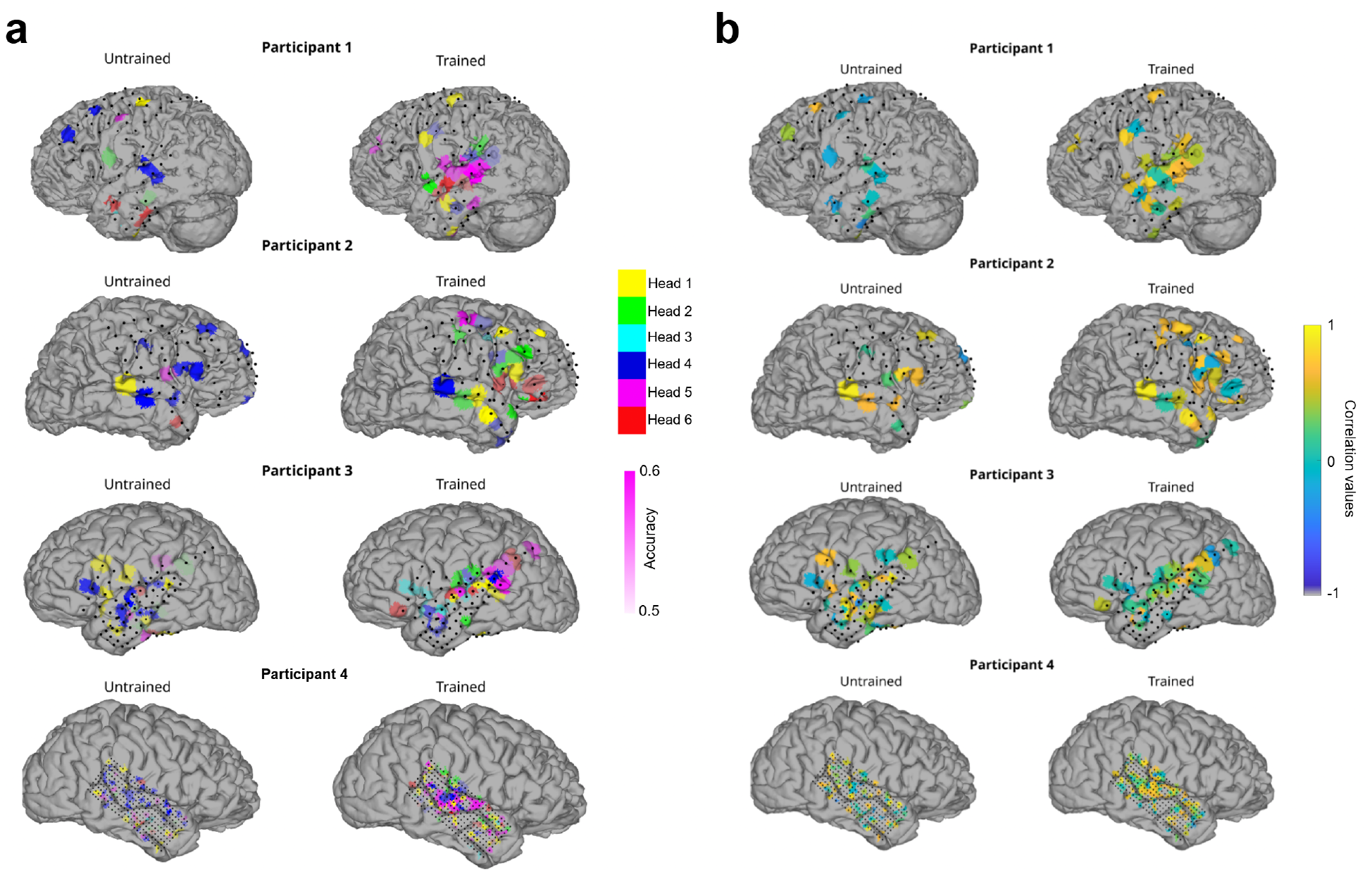
(**a**) Most predictive head for each significant electrode for the untrained and trained cortical MHSA network for 4 participants. The accuracy of the prediction is shown as transparency, scaled between 0.5 (chance level) and 0.6. Participant 3 is also shown in (4c). (**b**) Pairwise correlation was calculated between fold accuracies of the three-fold cross-validation for each contact independently and for each participant. The average pairwise correlation across the three possible fold pairs is plotted. High positive values indicate good consistency across the fold of the classification task.

### S2 Linear self-attention

Let ***x***_1_, …, ***x***_*T*_ be a sequence of *T* elements or tokens, with ***x***_*t*_ ∈ ℝ^*d*^. The self-attention mechanism central in transformer networks relies on three linear projections of sequence elements

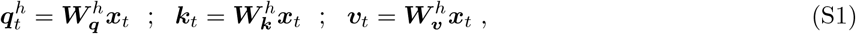

with 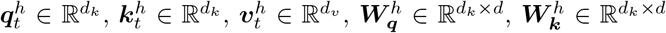, and 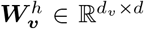. A classical self-attention mechanism (e.g. [4]) uses the softmax, or normalized exponential dot-product, as a similarity measure. It outputs the weighted sum across the *T* ^*h*^ past time steps,

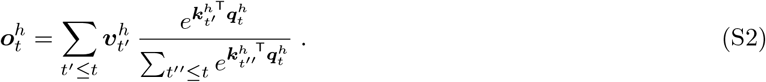

In this work, we instead use the alternative linear self-attention formulation, replacing the exponential kernel with a linear product of feature maps 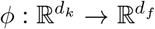 [36, 121]. Multiple works have shown how to approximate eq. S2 arbitrarily closely using projections to random features [122, 123]. We also introduce a memory discount factor 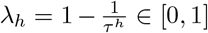 emerging from an effective integration time constant *τ* ^*h*^, corresponding to the memory length 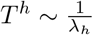 in eq. S2. The outputs are then a sum of discounted and (self-)attention-weighted values,

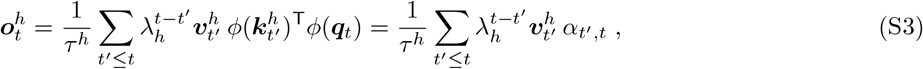

with attention 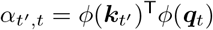 devoted at time *t* to the memory item from time *t*′, and suppressing the head index *h*. One advantage of non-normalized output (eq. S3) over the softmax-normalized output (eq. S2) is that, without normalization, the terms can be split into the product of a memory times the query,

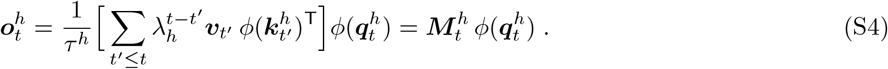

In the last equation, we introduced the key-value memory 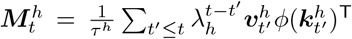, that can also be recurrently defined as in eq. 1.

An intuition for the memory parametrization in eq. 1 is obtained from the discretization of the continuous dynamics 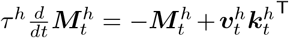, with a time step that either is set to unity as above, Δ*t* = 1, or can become infinitesimal, Δ*t* → *dt*. Eq. 1 is obtained from rearranging the terms of the time discrete approximation of this dynamics,

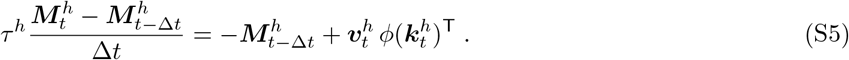

The linear transformer formulation without memory decay, i.e., with *λ*_*h*_ = 1 [38], is obtained back from setting the leak term on the right-hand side of eq. S5 to 0, while choosing *τ* ^*h*^ = Δ*t* = 1,

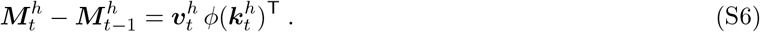

To obtain eq. 2 we first calculate from eq. 1 the explicit expression for ***M***_*t*_. Starting with ***M***_0_ = 0 at initial time *t* = 0, and proceeding for time steps *t*′ =Δ*t*, 2Δ*t*, 3Δ*t*, …, we get from iterating eq. 1,

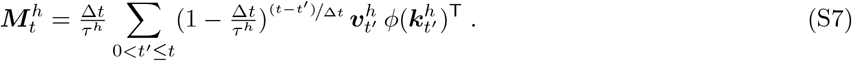

From this we get

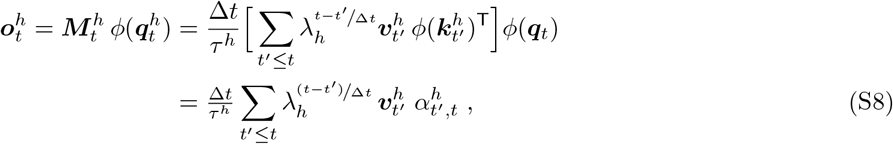

with 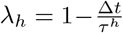, attention factor 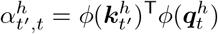, and the sum extending across *t*′ = *t, t* − Δ*t, t* − 2Δ*t*, … . For Δ*t* = 50 ms and *τ* ^*h*^ = 1 s, the values used in our Tiny Shakespeare simulations (figs. 2 and 3) and the translation task (fig. 5), we obtain the discount factor *λ*_*h*_ = 0.95.

The classical formulation of eq. S2 bears a computational cost that scales quadratically in time (sequence length), since each new query is compared with all the previous keys. In contrast, the rearrangement in eq. S4 only scales linearly in time, since each key-value pairing (outer product) can be computed only once, stored in a key-value memory, and reused for every query [36].

### S3 Stacking value-key-query codings from neurons to synapses to genes

Biological memories are distributed in space, e.g. across a hippocampal episodic memory and a cortical declarative memory, which allows the memories to bootstrap one another. This is achieved by sequentially stacking them, so that during the writing into the first memory the second memory is recruited, and the new item together with the second memory are jointly written into the first memory (see eq. 11). We may bootstrap a single memory with itself, formally by adding to the value in our L2/3 encoding the retrieved output of the hippocampal memory (suppressing the *h*-index),

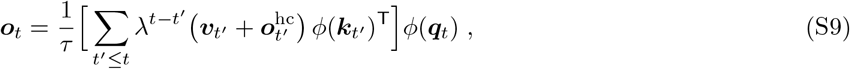

where in eq. S4 we added to ***v***_*t*_*′* the expression ***o***_*t*_*′* from eq. S3, evaluated at past time *t*′ so that values ***v***_*t*_*′′* enter in eq. S9 for times *t*^*′′*^ ≤ *t*′ ≤ *t*. We suppressed the RPE-scaling factor 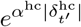 from eq. 11 for conceptual clarity.

The bracket in eq. S9 has a remarkable asymmetry: it combines the value from the neuronal memory with the output from the synaptic memory. But the synaptic memory 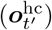, too, contains values, namely 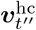 for times *t*^*′′*^ ≤ *t*′ further back in the past. These hippocampal values enter as an outer product 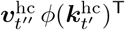 in the Hebbian update rule of the recurrent CA3 synapses, see e.g. [37]. As part of the induction of synaptic plasticity, 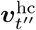 may itself be complemented by some protein kinase that determines the long-term potentiation [124]. Hence we may add to the hippocampal value a genetic readout, 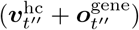, in the same way as we were adding to the L2/3 value the hippocampal readout, 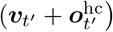, in eq. S9.

Hence, we may stack the value-key-query principle across different biological encoding levels. At the genetic level, the value-key-query principle is rather known as content-lock-probe principle. In fact, when reading out the genetic coding sequence of a protein (the mRNA receptor coding sequence, the content or value, ***v***_*t*_*′′* ), that coding sequence is flanked by a promotor sequence (CRE, the lock or key, ***k***_*t*_*′′* ) and it can be activated by a corresponding transcription factor (pCREB, the probe or query, ***k***_*t*_*′′*, [125]). It is remarkable that with three iterations of this content-lock-probe principle we span the breadth from neurons to genes.

### S4 Multi-head linear self-attention

The preceding subsections describe the computation in a single head of linear self-attention. We now extend to *n*_*h*_ heads. The classical presentation of multi-head self-attention proposes to concatenate the outputs 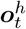 of the different heads and multiplies the resulting vector by output weight 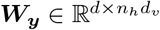, yielding the output

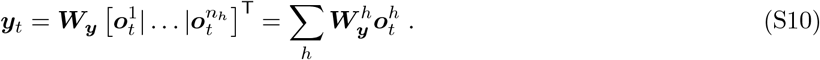

The last equation highlights attention heads as independent and additively integrated with head-specific output weights 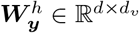.

Overall, the output ***y***_*t*_ in eq. S10 can be written as a sum of gain-modulated L5IT pyramidal neurons, summed up in L5ET output neurons in rows, and weighted by the output synapses,

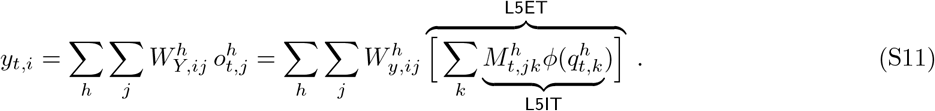

Finally, rather than having a single time constant *τ*, we opt to make time constants *τ* ^*h*^ head-specific, to allow each head to integrate information at different temporal rates (and to be selected based on the expected discounted future reward for each head/area, see eq. 9).

### S5 Formal gradients for the self-attention parameters

As we show, the explicit memory representation by our L2/3 neurons also allows for reassembling the appropriate learning signals required to implement gradient plasticity for the query- and readout-synapses. We derive the gradients of multi-head self-attention with respect to 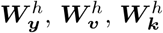, and 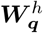 of a mean squared error loss in a self-supervised, autoregressive setting. Formally, the cost is defined as a squared mismatch between the output ***y***_*t*_ and the next input ***x***_*t*+1_ that should be prospectively predicted by the self-attention network,

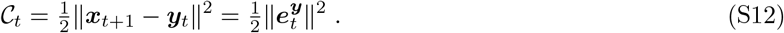

with ***y***_*t*_ defined by eq. 3. Such a cost function based on prospective prediction errors was recently suggested in the form of a least action principle for cortical computation [34], along with a dendritic implementation of the emerging learning rules [33, 31].

The formal derivatives of *C*_*t*_ with respect to parameters are (for Δ*t* = 1)

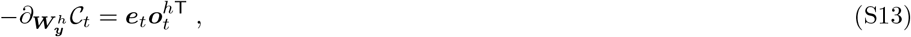

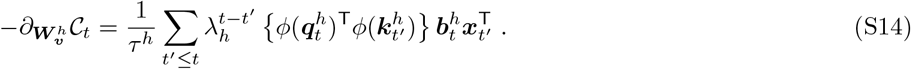

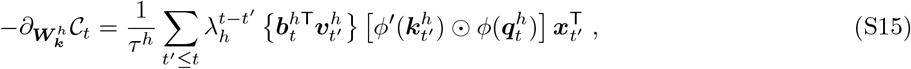

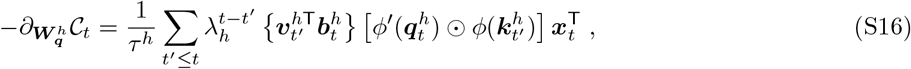

with 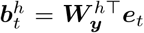 the backpropagated error, and 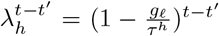 a scalar discounting factor. We checked these results against numerical approximations based on finite differences and forward automatic differentiation^1^. These gradients consist of outer products, scaled by scalars and a scalar product in the case of eqs. S14 to S16. Each component of the gradient matrices consist of a product of pre- and post-synaptic quantities, respecting the notion of locality.

The temporal summations over past tokens indicate slow variables influencing synaptic plasticity, stored and incremented with the presentation of each new token. We can write eqs. S14 to S16 in a recurrent form (using that the scalar products in the curly brackets {·} yield numbers, and hence commute with the corresponding subsequent column vector). For linear ‘dendritic’ transfer functions, *ϕ*(*x*) = *x* (such as in [126, 127], but generalizable for nonlinear *ϕ*, see below), we get

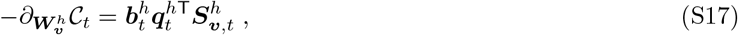

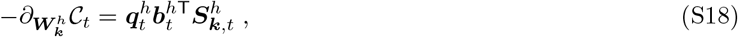

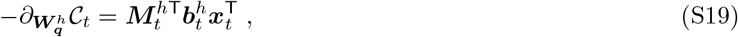

with matrix-valued slow synaptic variables 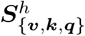 being recurrently updated according to

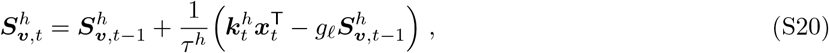

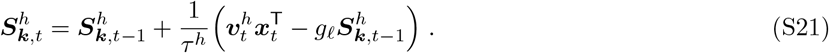

We next consider the general case with a nonlinear *ϕ* and Δ*t* = 1. Since the scalar 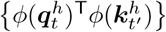 commutes with 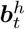, we can rewrite eq. S14 as

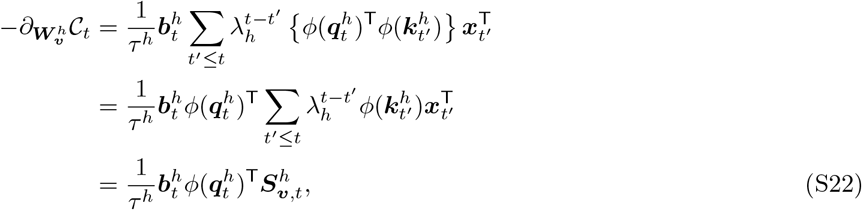

where

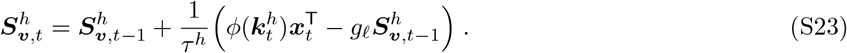

We can rewrite eq. S15 as

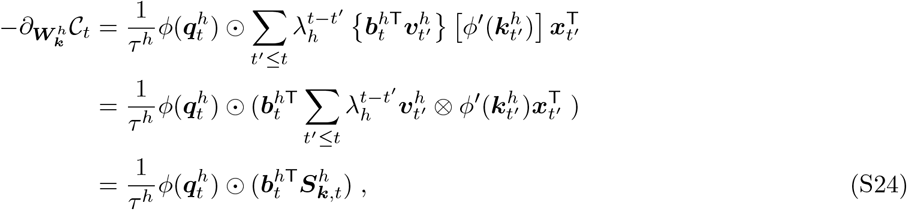

where

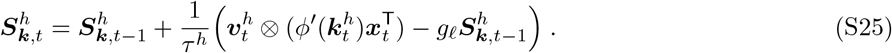

In the above, 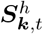 is a third-order tensor. We can rewrite eq. S16 as

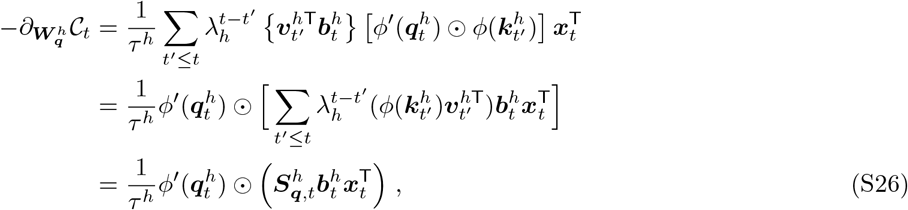

where

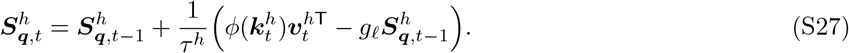

Notice that 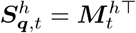, so that eq. S26 translates to the online rule 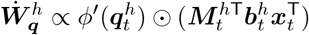. For *ϕ*(*x*) = *x*, eqs. S22, S24 and S26 simplify to eqs. S17, S18 and S19, respectively.

#### Local plasticity rules

The above energy gradients lead to the gradient-based plasticity of the corresponding synaptic weights from a presynaptic neuron *j* to a postsynaptic neuron *i*. For the readout and query-encoding synapses, the rules read as

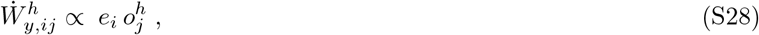

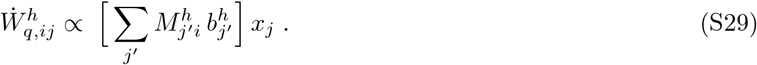

All the quantities are instantaneous function of time (for notational convenience we skipped the index *t*). The formulas generalize to non-linear *ϕ* where additional factors *ϕ*′ enter, without changing the overall structure.

#### Backpropagation across multiple cortico-thalamic loops

Here we consider the biological form of error backpropagation [31, 32] from the higher thalamic nucleus (encoding ***y***_*t*_) back to the cortical areas *X*^*h*^ and from these back to the sensory thalamic nucleus (encoding ***x***_*t*_). Following the notation in fig. 6b, the error in the higher thalamic nucleus is 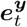, and the one in the sensory nucleus is 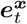. We consider the backpropagated error 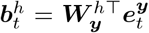, backpropagated via projection 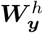 into area *X*^*h*^ and via projection 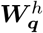 back into the thalamic kernel encoding ***x***_*t*_. We are interested in the plasticity rule for the readout synapses 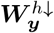 from a yet earlier cortical area *X*^*h*↓^, indexed by *h*↓, to that sensory nucleus ***x***_*t*_. In summary, the forward and backward information flow is

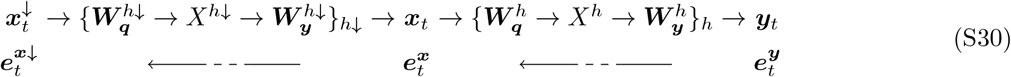

and we want to express the error 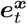 in the lower thalamic nucleus as a function of the error 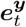 in the higher thalamic nucleus. Once knowing how to propagate from one kernel to the previous one (via 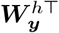 and 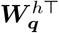, see eqs. S28 and S29), we can iterate the procedure to backpropagate across arbitrary deep self-attention networks.

For simplicity we consider a linear transfer function, *ϕ* = id, such that, following eq. S30 we have 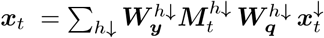 and 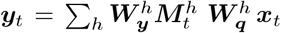. When backpropagating the error 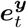, we have to back-propagate through the areas {*X*^*h*^}_*h*_, and this involves backpropagating through the memory matrices 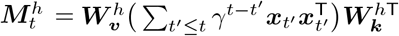.

The cost for the mapping across 2 thalamic loops is 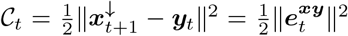, with 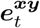 representing the error in predicting the next input 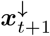 by the output ***y***_*t*_ two self-attention blocks further down. For simplicity we set Δ*t* = 1. We want to calculate the gradient of C_*t*_ with respect to 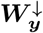 that projects out of the cortical areas {*X*^*h*↓^}_*h*↓_ into the thalamic nucleus ***x***_*t*_. We then calculate

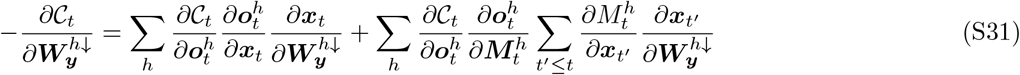

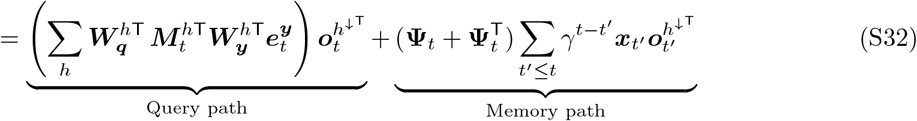

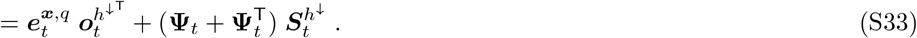

Here, 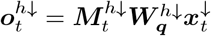 is the output from the upstream block, 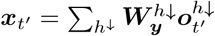 is the representation in the middle thalamic kernel (eq. S30), and 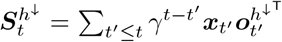 is the presynaptic eligibility trace [128, 129], a low-pass filter of pre- and postsynaptic coincidence with the same discount factor *γ* used for writing the key-value memory, updated online as 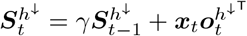.

The instantaneous ‘query error’ and the instantaneous ‘memory error’ in the lower thalamic kernel are read out from eq. S33 to be

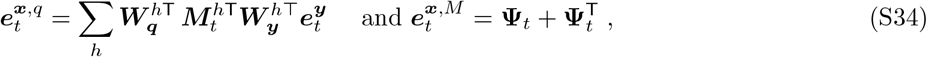

with contribution 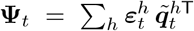 to the memory error, where 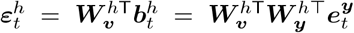 and 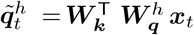 is the error and activity, respectively, backpropagated to the memory. The memory error contribution can alternatively be written as 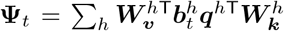, with 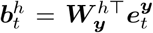. The individual query and memory errors, 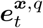 and 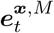, are driving the local plasticity when multiplied with the presynaptic activity 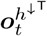 and the presynaptic eligibility trace 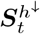 as expressed in eq. S33.

Both instantaneous errors, the query and the memory error (eq. S34), are constructed from the single instantaneous error 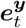 in the upper nucleus, and both are constructed through the same backward projection, 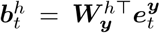. It is therefore enough to pass a single error between kernels and to reconstruct the pair of plasticity-driving errors locally at each stage. Since the memory error 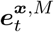 is a matrix, it cannot itself be represented as a propagated neuronal activity vector; for the propagation it is applied to the currently represented activity ***x***_*t*_. The total instantaneous error at the lower nucleus is then the vector

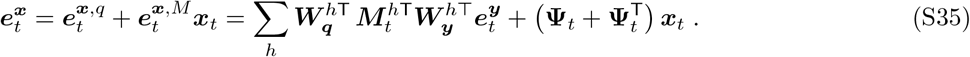

The next-lower kernel, representing 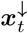 (see eq. S30), treats 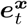 exactly as the upper kernel treated 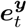. Through the backward projections 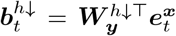 it locally reconstructs its own query and memory errors, 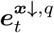 and 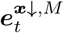 (i.e. eq. S34 one level down), which drive the local plasticity when multiplied with the presynaptic activity 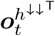 and the presynaptic eligibility trace 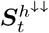, analogously to eq. S33. Applying the memory error to 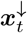 as in eq. S35 yields the total error 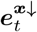 that is passed further back.

The scheme implements strict gradient descent for all synapses of the loop where the error arises (eqs. S28 and S29) and, owing to the eligibility trace, also for the readout synapses 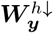 one loop further down. Hence, eq. S33 is the exact gradient of *C*_*t*_. For the query synapses of that loop, and for all synapses of yet deeper loops, the single propagated error retains the gradient components in which the error flows through the current retrieval operations and through the storage events whose temporal credit is resolved by the eligibility trace at the stage where the storage occurred. What is dropped, however, are ‘memory-through-memory’ components, in which the error would have to be delivered to network states ***x***_*t*_*′* at earlier times *t*′ *< t* (with weight 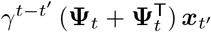 and travel from there through yet another, earlier memory.

### S6 Duality between inference and learning

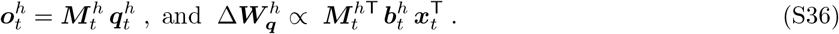

The duality statement reflects the fact that inference and plasticity involve a matrix 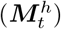 and its transpose 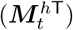,

The backpropagated error is 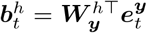, and that the inference step consists of calculating 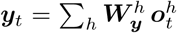. The learning rule is derived in eq. S29, and is identical to 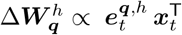 given in eq. 13, with 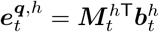. The duality at the level of 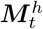 is also expressed in the symmetry between the L5 inference process and the L6 error calculation fig. 6b. It predicts a specific projection pattern of L3/3 pyramidal neurons to the apical dendrites of L5IT and L6IT pyramidal neurons engaging 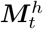 and its transpose, respectively.

### S7 MHSA pseudocodes annotated with biological substrates

#### Algorithm 1

Multi-head linear self-attention in cortico-thalamic circuits — One Layer, overview

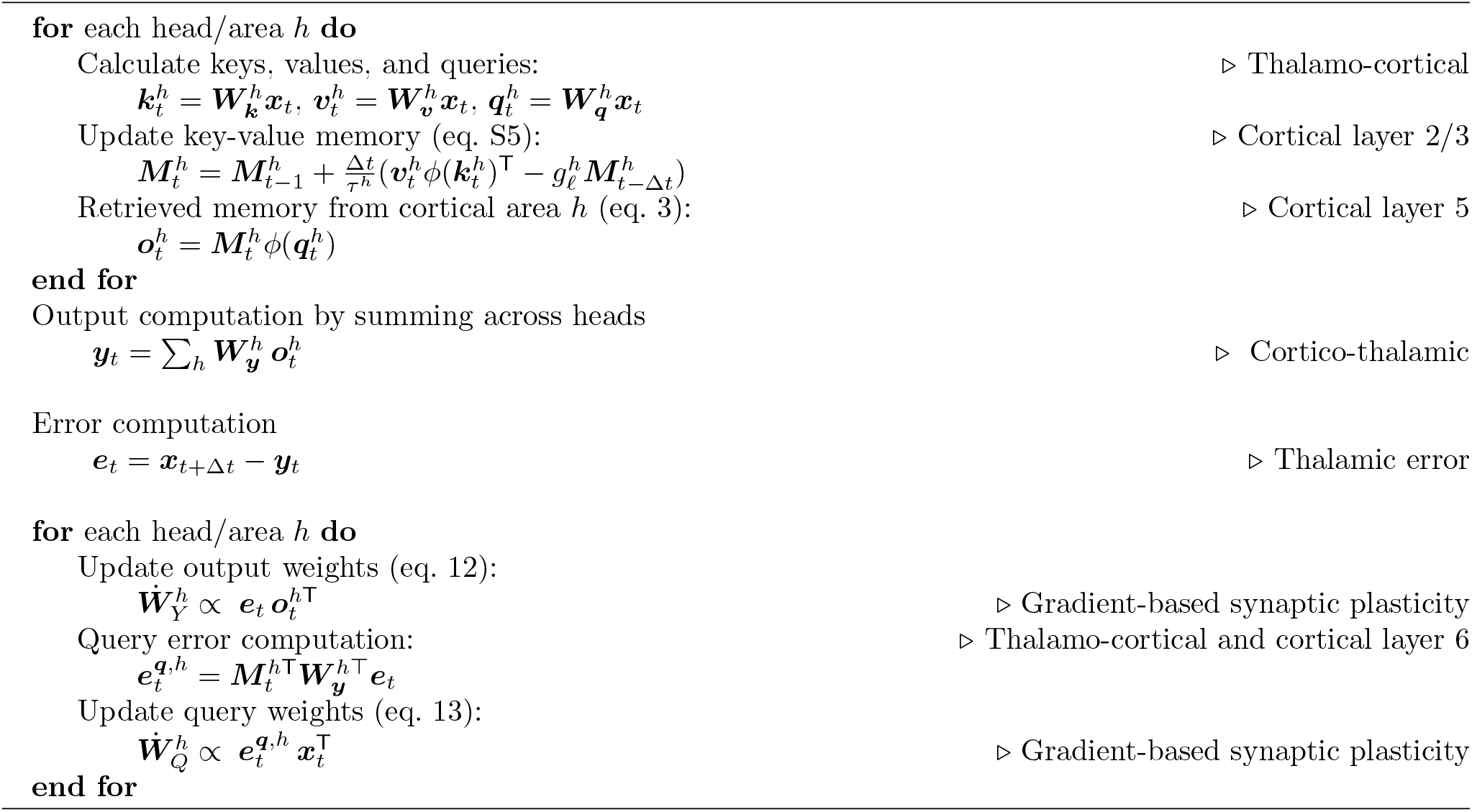

#### Algorithm 2

Multi-head linear Self-Attention in cortico-thalamic circuits — One layer, Details

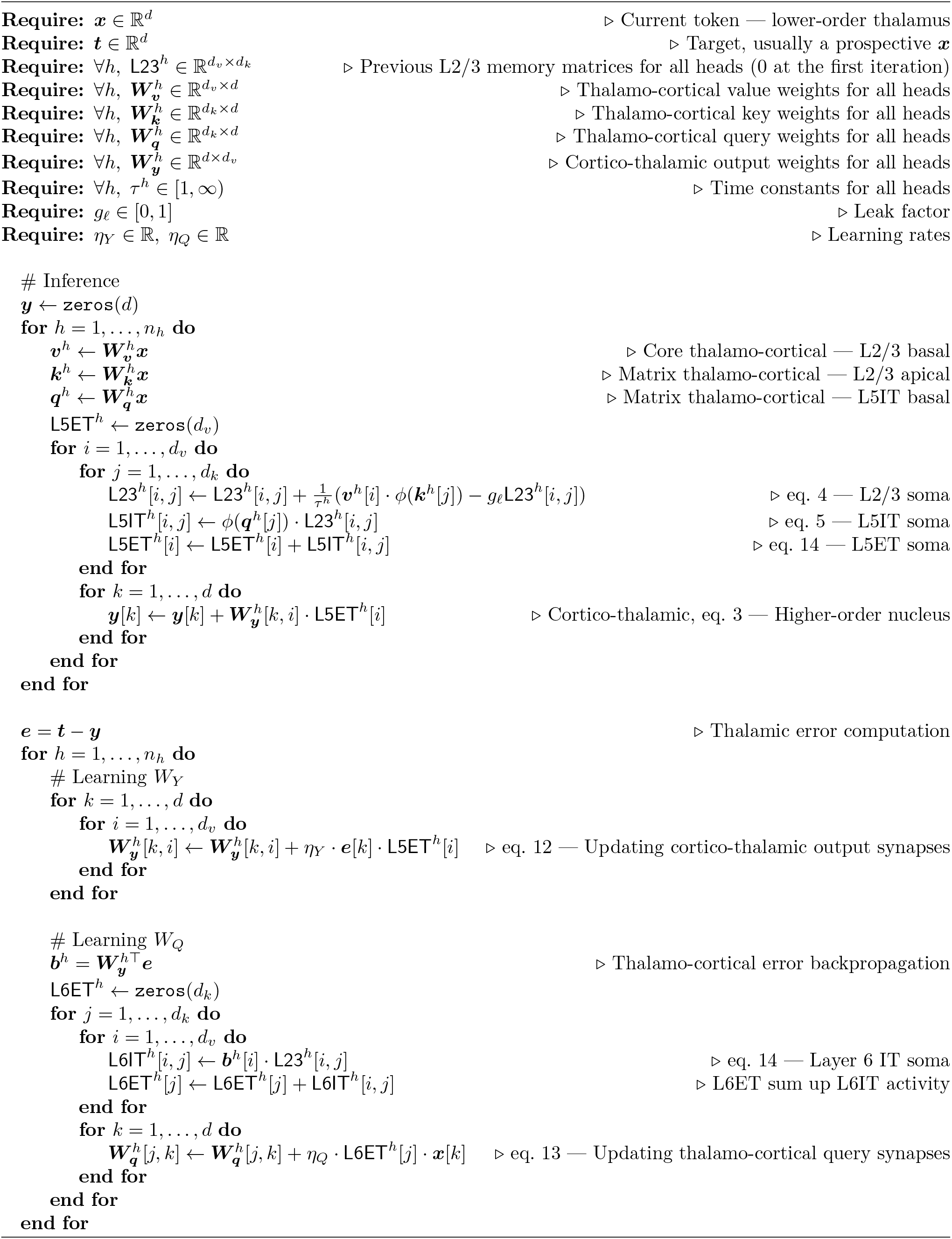

#### Algorithm 3

Multi-head linear self-attention in cortico-thalamic circuits — Inference & Learning

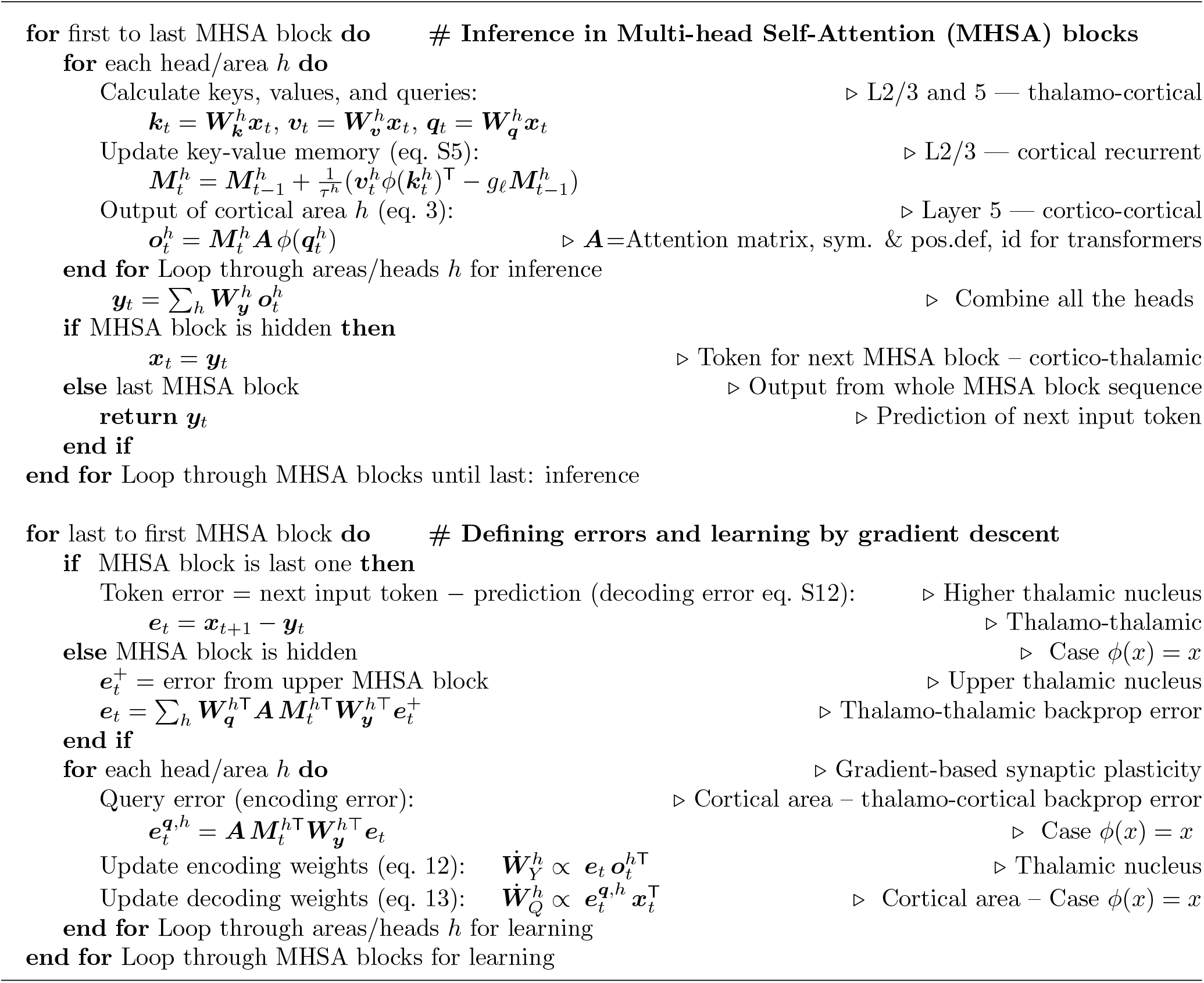

#### Algorithm 4

An equivalent formulation for 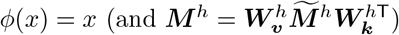 Inference and Plasticity

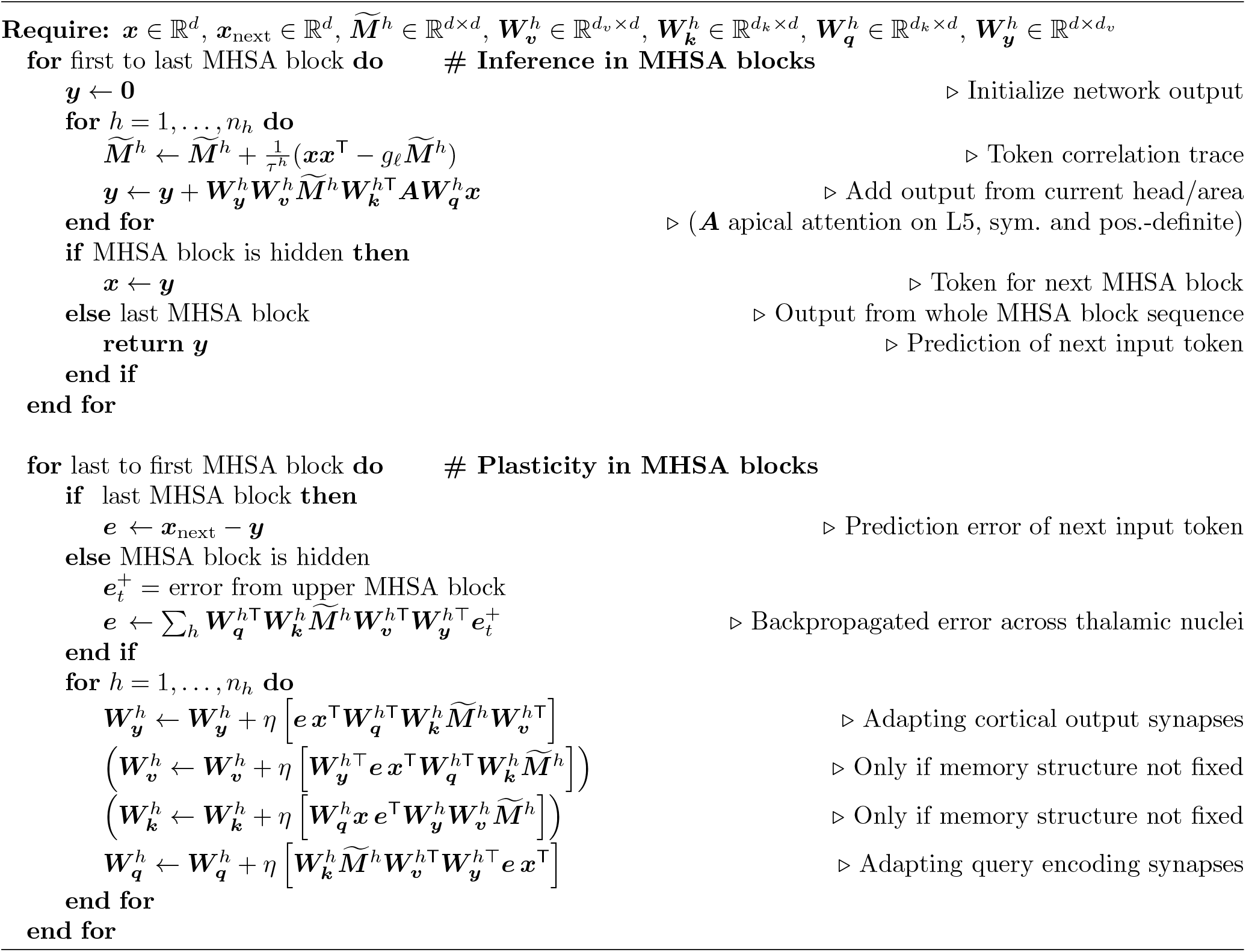

## Footnotes

1 https://github.com/arnogranier/Formal-MHSA-gradient-numerics

## References

[1] Viktor Jirsa, Huifang Wang, Paul Triebkorn, Meysam Hashemi, Jayant Jha, Jorge Gonzalez-martinez, Maxime Guye, and Julia Makhalova. “Personalised virtual brain models in epilepsy”. In: The Lancet Neurology 22.5 (2023), pp. 443–454. issn: 1474-4422. DOI: 10.1016/S1474-4422(23)00008-X. url: http://dx.doi.org/10.1016/S1474-4422(23)00008-X.

[2] Huifang E Wang, Paul Triebkorn, Martin Breyton, Borana Dollomaja, Jean-didier Lemarechal, Spase Petkoski, Pierpaolo Sorrentino, Damien Depannemaecker, Meysam Hashemi, and Viktor K Jirsa. “Virtual brain twins: from basic neuroscience to clinical use”. In: National Science Review 11 (2024). issn: 2053-714X. DOI: 10.1093/nsr/nwae079. url: https://doi.org/10.1093/nsr/nwae079.

[3] Meysam Hashemi, Damien Depannemaecker, Marisa Saggio, Paul Triebkorn, Giovanni Rabuffo, Jan Fousek, Abolfazl Ziaeemehr, Viktor Sip, Anastasios Athanasiadis, Martin Breyton, Marmaduke Woodman, Huifang Wang, Spase Petkoski, and Pierpaolo Sorrentino. “Principles and Operation of Virtual Brain Twins”. In: IEEE Reviews in Biomedical Engineering 19 (2026), pp. 111–139. DOI: 10.1109/RBME.2025.3562951.

[4] Ashish Vaswani, Noam Shazeer, Niki Parmar, Jakob Uszkoreit, Llion Jones, Aidan N Gomez, Lukasz Kaiser, and Illia Polosukhin. “Attention is all you need”. In: Advances in Neural Information Processing Systems. Vol. 30. 2017.

[5] Francois Chollet. “OpenAI o3 Breakthrough High Score on ARC-AGI-Pub”. In: ARC Prize Foundation (2024).

[6] Alexey Dosovitskiy, Lucas Beyer, Alexander Kolesnikov, Dirk Weissenborn, Xiaohua Zhai, Thomas Unterthiner, Mostafa Dehghani, Matthias Minderer, Georg Heigold, Sylvain Gelly, Jakob Uszkoreit, and Neil Houlsby. “An image is worth 16×16 words: transformers for image recognition at scale”. In: International Conference on Learning Representations. 2021.

[7] Kenneth D Harris and Gordon MG Shepherd. “The neocortical circuit: themes and variations”. In: Nature neuroscience 18.2 (2015), pp. 170–181.

[8] Nathaniel J Powell, Bettina Hein, Deyue Kong, Jonas Elpelt, Haleigh N Mulholland, Matthias Kaschube, and Gordon B Smith. “Common modular architecture across diverse cortical areas in early development”. In: Proceedings of the National Academy of Sciences 121.11 (2024), e2313743121.

[9] Emily E Meyer, Marcelina Martynek, Sabine Kastner, Margaret S Livingstone, and Michael J Arcaro. “Expansion of a conserved architecture drives the evolution of the primate visual cortex”. In: Proceedings of the National Academy of Sciences 122.3 (2025), e2421585122.

[10] Daniel J Felleman and David C Van Essen. “Distributed hierarchical processing in the primate cerebral cortex.” In: Cerebral cortex (New York, NY: 1991) 1.1 (1991), pp. 1–47.

[11] Rinaldo D D’Souza, Quanxin Wang, Weiqing Ji, Andrew M Meier, Henry Kennedy, Kenneth Knoblauch, and Andreas Burkhalter. “Hierarchical and nonhierarchical features of the mouse visual cortical network”. In: Nature communications 13.1 (2022), p. 503.

[12] Jochen F Staiger and Carl CH Petersen. “Neuronal circuits in barrel cortex for whisker sensory perception”. In: Physiological reviews 101.1 (2021), pp. 353–415.

[13] Meike Sievers, Alessandro Motta, Martin Schmidt, Yagmur Yener, Sahil Loomba, Kun Song, Johannes Bruett, and Moritz Helmstaedter. “Connectomic reconstruction of a cortical column”. In: bioRxiv (2024), pp. 2024–03.

[14] The MICrONS consortium. “Functional connectomics spanning multiple areas of mouse visual cortex”. In: Nature 640.8058 (2025), pp. 435–447.

[15] Kazuyuki Samejima, Yasumasa Ueda, and Kenji Doya. “Representation of Action-Specific Reward Values in the Striatum”. In: Science 310.November (2005), pp. 1337–1341.

[16] Mitsuko Watabe-Uchida, Neir Eshel, and Naoshige Uchida. “Neural Circuitry of Reward Prediction Error”. In: Annual Review of Neuroscience 40 (2017), pp. 373–394.

[17] Travis D Goode, Kazumasa Z Tanaka, Amar Sahay, and Thomas J Mchugh. “An Integrated Index: Engrams, Place Cells, and Hippocampal Memory”. In: Neuron 107.5 (2020), pp. 805–820. issn: 0896-6273. DOI: 10.1016/j.neuron.2020.07.011. url: https://doi.org/10.1016/j.neuron.2020.07.011.

[18] Ji Hye Lee, Woong Bin Kim, Eui Ho Park, and Jun Hyeong Cho. “Neocortical synaptic engrams for remote contextual memories”. In: Nature Neuroscience 26.2 (2023), pp. 259–273. issn: 15461726. DOI: 10.1038/s41593-022-01223-1.

[19] Shohei Furutachi, Alexis D Franklin, Andreea M Aldea, Thomas D Mrsic-Flogel, and Sonja B Hofer. “Cooperative thalamocortical circuit mechanism for sensory prediction errors”. In: Nature 633.8029 (2024), pp. 398–406.

[20] Claire McKinnon, Christina Mo, and S Murray Sherman. “Disruption of Transthalamic Circuitry from the Primary Visual Cortex Impairs Visual Discrimination in Mice”. In: Journal of Neuroscience 45.18 (2025).

[21] Christopher J Whyte, Eli J Müller, Jaan Aru, Matthew Larkum, Yohan John, Brandon R Munn, and James M Shine. “A burst-dependent thalamocortical substrate for perceptual awareness”. In: PLOS Computational Biology 21.4 (2025), e1012951.

[22] Federico Benitez, Cyriel Pennartz, and Walter Senn. “The conductor model of consciousness, our neuromorphic twins, and the human-AI deal”. In: AI and Ethics (2024), pp. 1–23. url: https://doi.org/10.1007/s43681-024-00580-w.

[23] Gwendolin Schoenfeld, Sepp Kollmorgen, Matthias C Tsai, Christopher Lewis, Shuting Han, Philipp Bethge, Anna Maria Reuss, Adriano Aguzzi, and Walter Senn. “Unsigned temporal difference errors in cortical L5 dendrites during learning”. In: bioRxiv (2024), pp. 1–42.

[24] Valerio Francioni, Vincent D Tang, Enrique H S Toloza, Zilan Ding, Norma J Brown, and Mark T Harnett. “Vectorized instructive signals in cortical dendrites”. In: Nature 652.April (2026), pp. 1254–1263. issn: 1476-4687. DOI: 10.1038/s41586-026-10190-7. url: http://dx.doi.org/10.1038/s41586-026-10190-7.

[25] James C. R. Whittington, Joseph Warren, and Tim E.J. Behrens. “Relating transformers to models and neural representations of the hippocampal formation”. In: International Conference on Learning Representations. 2022.

[26] Leo Kozachkov, Ksenia V Kastanenka, and Dmitry Krotov. “Building transformers from neurons and astrocytes”. In: Proceedings of the National Academy of Sciences 120.34 (2023), e2219150120.

[27] Arno Granier and Walter Senn. “Multihead self-attention in cortico-thalamic circuits”. In: arxiv 2 (2025), pp. 1–15. arXiv: arXiv:2504.06354v3. url: https://arxiv.org/pdf/2504.06354.

[28] Peter König and Mario Negrello. “The neuroscience of transformers”. In: arXiv (2026), pp. 1–34. arXiv: arXiv:2603.15339v1.

[29] Hubert Ramsauer, Bernhard Schäfl, Johannes Lehner, Philipp Seidl, Michael Widrich, Lukas Gruber, Markus Holzleitner, Thomas Adler, David Kreil, Michael K Kopp, Günter Klambauer, Johannes Brandstetter, and Sepp Hochreiter. “Hopfield Networks is All You Need”. In: International Conference on Learning Representations. 2021.

[30] Dmitry Krotov and John J. Hopfield. “Large Associative Memory Problem in Neurobiology and Machine Learning”. In: International Conference on Learning Representations. 2021.

[31] João Sacramento, Yoshua Bengio, Rui Ponte Costa, and Walter Senn. “Dendritic cortical microcircuits approximate the backpropagation algorithm”. In: Advances in Neural Information Processing Systems (NeurIPS) 2018 (2018), pp. 8721–8732. issn: 10495258. arXiv: 1810.11393. url: https://intranet.physio.unibe.ch/Publikationen/Dokumente/Sacramento2018Dendriticc.pdf.

[32] Kevin Max, Laura Kriener, Garibaldi Pineda García, Thomas Nowotny, Ismael Jaras, Walter Senn, and Mihai A. Petrovici. “Learning efficient backprojections across cortical hierarchies in real time”. In: Nature Machine Intelligence 6.6 (2024), pp. 619–630. issn: 25225839. DOI: 10.1038/s42256-024-00845-3. arXiv: 2212.10249. url: http://dx.doi.org/10.1038/s42256-024-00845-3.

[33] Robert Urbanczik and Walter Senn. “Learning by the dendritic prediction of somatic spiking”. In: Neuron 81.3 (2014), pp. 521–528.

[34] Walter Senn, Dominik Dold, Akos F Kungl, Benjamin Ellenberger, Jakob Jordan, Yoshua Bengio, João Sacramento, and Mihai A Petrovici. “A neuronal least-action principle for real-time learning in cortical circuits”. In: ELife 12 (2024), RP89674.

[35] Teuvo Kohonen. “Correlationn Matrix Memories”. In: IEEE Transactions on Computers c-21.4 (1972), pp. 353–359.

[36] Angelos Katharopoulos, Apoorv Vyas, Nikolaos Pappas, and François Fleuret. “Transformers are RNNs: Fast autoregressive transformers with linear attention”. In: International conference on machine learning. PMLR. 2020, pp. 5156–5165.

[37] Samuel J Gershman, Ila Fiete, and Kazuki Irie. “Key-value memory in the brain”. In: Neuron 113.11 (2025), pp. 1694–1707.

[38] Virginie van Wassenhove, Ken W. Grant, and David Poeppel. “Temporal window of integration in auditory-visual speech perception”. In: Neuropsychologia 45.3 (2007), pp. 598–607. issn: 00283932. DOI: 10.1016/j.neuropsychologia.2006.01.001.

[39] Elodie Fino and Rafael Yuste. “Dense inhibitory connectivity in neocortex”. In: Neuron 69.6 (2011), pp. 1188–1203.

[40] Alison L Barth and James FA Poulet. “Experimental evidence for sparse firing in the neocortex”. In: Trends in neurosciences 35.6 (2012), pp. 345–355.

[41] Jean-Sébastien Jouhanneau, Jens Kremkow, and James FA Poulet. “Single synaptic inputs drive high-precision action potentials in parvalbumin expressing GABA-ergic cortical neurons in vivo”. In: Nature communications 9.1 (2018), p. 1540.

[42] Carl Holmgren, Tibor Harkany, Björn Svennenfors, and Yuri Zilberter. “Pyramidal cell communication within local networks in layer 2/3 of rat neocortex”. In: Journal of Physiology 551.1 (2003), pp. 139–153. issn: 00223751. DOI: 10.1113/jphysiol.2003.044784.

[43] Kenta Funayama, Genki Minamisawa, Nobuyoshi Matsumoto, Hiroshi Ban, Allen W. Chan, Norio Matsuki, Timothy H. Murphy, and Yuji Ikegaya. “Neocortical rebound depolarization enhances visual perception”. In: PLoS Biology 13.8 (2015), pp. 1–25. issn: 15457885. DOI: 10.1371/journal.pbio.1002231.

[44] Nelson Spruston. “Pyramidal neurons: dendritic structure and synaptic integration”. In: Nature Reviews Neuroscience 9.3 (2008), pp. 206–221.

[45] Matthew E Larkum, Walter Senn, and Hans-R Lüscher. “Top-down dendritic input increases the gain of layer 5 pyramidal neurons”. In: Cerebral cortex 14.10 (2004), pp. 1059–1070.

[46] Marcel Oberlaender, Zimbo S R M Boudewijns, Tatjana Kleele, Huibert D Mansvelder, and Bert Sakmann. “Three-dimensional axon morphologies of individual layer 5 neurons indicate cell type-specific intracortical pathways for whisker motion and touch”. In: Proc. Nat. Acad. Sci. (PNAS) USA 108.10 (2011). DOI: 10.1073/pnas.1100647108.

[47] Rodney J Douglas and Kevan AC Martin. “Neuronal circuits of the neocortex”. In: Annu. Rev. Neurosci. 27.1 (2004), pp. 419–451.

[48] Michael Quiquempoix, Sophie L Fayad, Katia Boutourlinsky, Nathalie Leresche, Régis C Lambert, and Thomas Bessaih. “Layer 2/3 pyramidal neurons control the gain of cortical output”. In: Cell reports 24.11 (2018), pp. 2799–2807.

[49] Leopoldo Petreanu, Tianyi Mao, Scott M Sternson, and K Svoboda. “The subcellular organization of neocortical excitatory connections.” In: Nature 457.7233 (2009), pp. 1142–5. issn: 1476-4687. DOI: 10.1038/nature07709. arXiv: NIHMS150003. url: http://www.pubmedcentral.nih.gov/articlerender.fcgi?artid=2745650{\&}tool=pmcentrez{\&}rendertype=abstract.

[50] Anton Sumser, Rebecca A Mease, Bert Sakmann, and Alexander Groh. “Organization and somatotopy of corticothalamic projections from L5B in mouse barrel cortex”. In: Proc. Nat. Acad. Sci. (PNAS) USA 114.33 (2017), pp. 8853–8858. DOI: 10.1073/pnas.1704302114.

[51] S Murray Sherman and W Martin Usrey. “Transthalamic Pathways for Cortical Function”. In: Journal of Neuroscience 44.35 (2024).

[52] EG Jones. “The core and matrix of thalamic organization”. In: Neuroscience 85.2 (1998), pp. 331–345.

[53] David Mumford. “On the computational architecture of the neocortex: I. The role of the thalamo-cortical loop”. In: Biological cybernetics 65.2 (1991), pp. 135–145.

[54] Edgar A DeYoe and David C Van Essen. “Concurrent processing streams in monkey visual cortex”. In: Trends in neurosciences 11.5 (1988), pp. 219–226.

[55] Vandana Sampathkumar, Andrew Miller-Hansen, S Murray Sherman, and Narayanan Kasthuri. “Integration of signals from different cortical areas in higher order thalamic neurons”. In: Proceedings of the National Academy of Sciences 118.30 (2021), e2104137118.

[56] S. Murray Sherman. “Thalamus plays a central role in ongoing cortical functioning”. In: Nature Neuroscience 19.4 (2016), pp. 533–541. issn: 15461726. DOI: 10.1038/nn.4269.

[57] Dmitry Krotov and John J. Hopfield. “Dense associative memory for pattern recognition”. In: Advances in Neural Information Processing Systems Nips (2016), pp. 1180–1188. issn: 10495258. arXiv: 1606.01164.

[58] Ali Rahimi and Benjamin Recht. “Random features for large-scale kernel machines”. In: Advances in neural information processing systems 20 (2007).

[59] Yihe Dong, Lorenzo Noci, Mikhail Khodak, and Mufan Li. “Is Random Attention Sufficient for Sequence Modeling? Disentangling Trainable Components in the Transformer”. In: arXiv preprint arXiv:2506.01115 (2025).

[60] Laura N. Driscoll, Lea Duncker, and Christopher D. Harvey. “Representational drift: Emerging theories for continual learning and experimental future directions”. In: Current Opinion in Neurobiology 76 (2022), p. 102609. issn: 18736882. DOI: 10.1016/j.conb.2022.102609. url: https://doi.org/10.1016/j.conb.2022.102609.

[61] Satoshi Manita, Takayuki Suzuki, Chihiro Homma, Takashi Matsumoto, Maya Odagawa, Kazuyuki Yamada, Keisuke Ota, Chie Matsubara, Ayumu Inutsuka, Masaaki Sato, Masamichi Ohkura, Akihiro Ya-manaka, Yuchio Yanagawa, Junichi Nakai, Yasunori Hayashi, Matthew E. Larkum, and Masanori Murayama. “A Top-Down Cortical Circuit for Accurate Sensory Perception”. In: Neuron 86.5 (2015), pp. 1–6. issn: 08966273. DOI: 10.1016/j.neuron.2015.05.006. url: http://linkinghub.elsevier.com/retrieve/pii/S0896627315004134.

[62] Mária Ercsey-Ravasz, Nikola T Markov, Camille Lamy, David C Van Essen, Kenneth Knoblauch, Zoltán Toroczkai, and Henry Kennedy. “A predictive network model of cerebral cortical connectivity based on a distance rule”. In: Neuron 80.1 (2013), pp. 184–197.

[63] Kenneth D Harris and Thomas D Mrsic-Flogel. “Cortical connectivity and sensory coding”. In: Nature 503.7474 (2013), pp. 51–58.

[64] Asif A Ghazanfar and Charles E Schroeder. “Is neocortex essentially multisensory?” In: Trends in cognitive sciences 10.6 (2006), pp. 278–285.

[65] Tianyi Mao, Deniz Kusefoglu, Bryan M Hooks, Daniel Huber, Leopoldo Petreanu, and Karel Svoboda. “Long-range neuronal circuits underlying the interaction between sensory and motor cortex”. In: neuron 72.1 (2011), pp. 111–123.

[66] Jan Kubanek and Gerwin Schalk. “NeuralAct: A Tool to Visualize Electrocortical (ECoG) Activity on a Three-Dimensional Model of the Cortex”. In: Neuroinformatics 13.2 (Apr. 2015), pp. 167–174. issn: 1559-0089. DOI: 10.1007/s12021-014-9252-3.

[67] Peter Hagoort. “The neurobiology of language beyond single-word processing”. In: Science 58.October (2019), pp. 55–58.

[68] Gregory Hickok and David Poeppel. “The cortical organization of speech processing”. In: Nature Reviews Neuroscience 8.May (2007), pp. 393–402.

[69] John P Aggleton, Shane M O Mara, Seralynne D Vann, Nick F Wright, Marian Tsanov, and Jonathan T Erichsen. “Hippocampal – anterior thalamic pathways for memory : uncovering a network of direct and indirect actions”. In: European Journal ofNeuroscience 31 (2010), pp. 2292–2307. DOI: 10.1111/j.1460-9568.2010.07251.x.

[70] Pierre Lavenex and David G Amaral. “Hippocampal-Neocortical Interaction: A Hierarchy of Associativity”. In: Hippocampus 430 (2000), pp. 420–430.

[71] Sara Moberg and Naoya Takahashi. “Neocortical layer 5 subclasses: From cellular properties to roles in behavior”. In: Frontiers in Synaptic Neuroscience 14.October (2022), pp. 1–12. issn: 16633563. DOI: 10.3389/fnsyn.2022.1006773.

[72] Kenji Doya. “Complementary roles of basal ganglia and cerebellum in learning and motor control”. In: Current Opinion in Neurobiology 10 (2000), pp. 732–739.

[73] Robert S. Turner and Michel Desmurget. “Basal ganglia contributions to motor control: A vigorous tutor”. In: Current Opinion in Neurobiology 20.6 (2010), pp. 704–716. issn: 09594388. DOI: 10.1016/j.conb.2010.08.022. url: http://dx.doi.org/10.1016/j.conb.2010.08.022.

[74] Arthur Leblois, Thomas Boraud, and David Hansel. “Reward-driven adaptation of movements requires strong recurrent basal ganglia – cortical loops”. In: PNAS 122.50 (2025), pp. 1–11. DOI: 10.1073/pnas. url: https://doi.org/10.1073/pnas.2515994122.

[75] Zengcai V. Guo, Hidehiko K. Inagaki, Kayvon Daie, Shaul Druckmann, Charles R. Gerfen, and Karel Svoboda. “Maintenance of persistent activity in a frontal thalamocortical loop”. In: Nature 545.7653 (2017), pp. 181–186. issn: 14764687. DOI: 10.1038/nature22324.

[76] Bryan M Hooks, Andrew E Papale, Ronald F Paletzki, Muhammad W Feroze, Brian S Eastwood, Jonathan J Couey, Johan Winnubst, Jayaram Chandrashekar, and Charles R Gerfen. “Topographic precision in sensory and motor corticostriatal projections varies across cell type and cortical area”. In: Nature communications 9.1 (2018), p. 3549.

[77] Ethan S. Bromberg-Martin, Masayuki Matsumoto, and Okihide Hikosaka. “Dopamine in Motivational Control: Rewarding, Aversive, and Alerting”. In: Neuron 68.5 (2010), pp. 815–834. issn: 08966273. DOI: 10.1016/j.neuron.2010.11.022. url: http://dx.doi.org/10.1016/j.neuron.2010.11.022.

[78] Wolfram Schultz, Peter Dayan, and P Read Montague. “A neural substrate of prediction and reward”. In: Science 275.5306 (1997), pp. 1593–1599.

[79] Masayuki Matsumoto and Okihide Hikosaka. “Two types of dopamine neuron distinctly convey positive and negative motivational signals”. In: Nature 459 (2009), pp. 837–842. issn: 0028-0836. DOI: 10.1038/nature08028.

[80] Matthias C Tsai, Jasper Teutsch, Willem A M Wybo, Fritjof Helmchen, Abhishek Banerjee, and Walter Senn. “Hierarchy of prediction errors shapes the learning of context-dependent sensory representations”. In: bioRxiv (2024), pp. 1–32.

[81] Johanni Brea, Alexisz Tamás Gaál, Robert Urbanczik, and Walter Senn. “Prospective Coding by Spiking Neurons”. In: PLoS Computational Biology 12.6 (2016), pp. 1–25. issn: 1553-7358. DOI: 10.1371/journal.pcbi.1005003. url: http://dx.plos.org/10.1371/journal.pcbi.1005003.

[82] Kishore Papineni, Salim Roukos, Todd Ward, and Wei-jing Zhu. “B LEU: a Method for Automatic Evaluation of Machine Translation”. In: Proceedings of the 40th Annual Meeting of the Association for Computational Linguistics (ACL) July (2002), pp. 311–318.

[83] Gwendolyn English, Newsha Ghasemi Nejad, Marcel Sommerfelt, Mehmet Fatih Yanik, and Wolfger Von der Behrens. “Bayesian surprise shapes neural responses in somatosensory cortical circuits”. In: Cell Reports 42.2 (2023).

[84] Rachel M Cassidy, Angel V Macias, Willian N Lagos, Chiamaka Ugorji, and Edward M Callaway. “Complementary organization of mouse driver and modulator cortico-thalamo-cortical circuits”. In: Journal of Neuroscience 45.5 (2025).

[85] Alejandro Tabas, Glad Mihai, Stefan Kiebel, and Robert Trampel. “Abstract rules drive adaptation in the subcortical sensory pathway”. In: eLife 9 (2020), pp. 1–19.

[86] Carmen Varela, Joao V S Moreira, Basak Kocaoglu, Salvador Dura-bernal, Subutai Ahmad, and Robert Worden. “A mechanism for deviance detection and contextual routing in the thalamus : a review and theoretical proposal”. In: Frontiers in Neuroscience February (2024), pp. 1–15. DOI: 10.3389/fnins.2024.1359180.

[87] Manuel A Castro-alamancos and Maria E Calcagnotto. “Presynaptic Long-Term Potentiation in Corticothalamic Synapses”. In: The Journal of Neuroscience 19.20 (1999), pp. 9090–9097.

[88] Jeremy S Biane, Yoshio Takashima, Massimo Scanziani, James M Conner, Mark H Tuszynski, Jeremy S Biane, Yoshio Takashima, Massimo Scanziani, James M Conner, and Mark H Tuszynski. “Thalamocortical projections onto behaviorally relevant neurons exhibit plasticity during adult motor learning”. In: Neuron 89.6 (2016), pp. 1173–1179. issn: 0896-6273. DOI: 10.1016/j.neuron.2016.02.001. url: http://dx.doi.org/10.1016/j.neuron.2016.02.001.

[89] Richard H.R. Hahnloser, Rahul Sarpeshkar, Misha A. Mahowald, Rodney J. Douglas, and H. Sebastian Seung. “Digital selection and analogue amplication coexist in a cortex-inspired silicon circuit”. In: Nature 442.1998 (2000), pp. 947–951.

[90] Masanori Murayama, Enrique Pérez-Garci, Thomas Nevian, Tobias Bock, Walter Senn, and Matthew E Larkum. “Dendritic encoding of sensory stimuli controlled by deep cortical interneurons.” In: Nature 457.February (2009), pp. 1137–1141. issn: 0028-0836. DOI: 10.1038/nature07663.

[91] Matteo Carandini and David J Heeger. “Normalization as a canonical neural computation”. In: Nature reviews neuroscience 13.1 (2012), pp. 51–62.

[92] Yang Shen, Julia Wang, and Saket Navlakha. “A correspondence between normalization strategies in artificial and biological neural networks”. In: Neural computation 33.12 (2021), pp. 3179–3203.

[93] Martin Schrimpf, Jonas Kubilius, Michael J Lee, N Apurva Ratan Murty, and Robert Ajemian. “Integrative Benchmarking to Advance Neurally Mechanistic Models of Human Intelligence”. In: Neuron 108.3 (2020), pp. 413–423. issn: 0896-6273. DOI: 10.1016/j.neuron.2020.07.040. url: https://doi.org/10.1016/j.neuron.2020.07.040.

[94] Abdulkadir Gokce and Martin Schrimpf. “Scaling Laws for Task-Optimized Models of the Primate Visual Ventral Stream”. In: Proceedings of the 42 nd International Conference on Machine Learning 267 (2025), pp. 1–22. arXiv: arXiv:2411.05712v3. url: https://arxiv.org/pdf/2411.05712.

[95] Henry Markram et al. “Reconstruction and Simulation of Neocortical Microcircuitry”. In: Cell 163.2 (2015), pp. 456–492. issn: 00928674. DOI: 10.1016/j.cell.2015.09.029. url: http://linkinghub.elsevier.com/retrieve/pii/S0092867415011915.

[96] Gaute Einevoll, Alain Destexhe, Markus Diesmann, Sonja Grün, Viktor Jirsa, Marc De Kamps, Michele Migliore, Torbjorn V. Ness, Hans Plesser E., and Felix Schürmann. “The Scientific Case for Brain Simulations”. In: Neuron 102 (2019), pp. 735–744. DOI: 10.1016/j.neuron.2019.03.027.

[97] Yazan N. Billeh, Binghuang Cai, Sergey L. Gratiy, Kael Dai, Ramakrishnan Iyer, Nathan W. Gouwens, Reza Abbasi-Asl, Xiaoxuan Jia, Joshua H. Siegle, Shawn R. Olsen, Christof Koch, Stefan Mihalas, and Anton Arkhipov. “Systematic Integration of Structural and Functional Data into Multi-scale Models of Mouse Primary Visual Cortex”. In: Neuron 106.3 (2020), 388–403.e18. issn: 10974199. DOI: 10.1016/j.neuron.2020.01.040. url: https://doi.org/10.1016/j.neuron.2020.01.040.

[98] Thomas B Christophel, P Christiaan Klink, Bernhard Spitzer, and Pieter R Roelfsema. “The Distributed Nature of Working Memory”. In: Trends in Cognitive Sciences 21.2 (2017), pp. 111–124. issn: 1364-6613. DOI: 10.1016/j.tics.2016.12.007. url: http://dx.doi.org/10.1016/j.tics.2016.12.007.

[99] Marcelo G Mattar and Nathaniel D Daw. “Prioritized memory access explains planning and hippocampal replay”. In: Nature Neuroscience 21.November (2018), pp. 1609–1617. issn: 1546-1726. DOI: 10.1038/s41593-018-0232-z. url: http://dx.doi.org/10.1038/s41593-018-0232-z.

[100] Joao Barbosa, Heike Stein, Rebecca L Martinez, Adrià Galan-gadea, Sihai Li, Josep Dalmau, Kirsten C S Adam, Josep Valls-solé, Christos Constantinidis, and Albert Compte. “Interplay between persistent activity and activity-silent dynamics in the prefrontal cortex underlies serial biases in working memory”. In: Nature Neuroscience 23.August (2020), pp. 16–18. issn: 1546-1726. DOI: 10.1038/s41593-020-0644-4. url: http://dx.doi.org/10.1038/s41593-020-0644-4.

[101] George A. Mashour, Pieter Roelfsema, Jean Pierre Changeux, and Stanislas Dehaene. “Conscious Processing and the Global Neuronal Workspace Hypothesis”. In: Neuron 105.5 (2020), pp. 776–798. issn: 10974199. DOI: 10.1016/j.neuron.2020.01.026. url: https://doi.org/10.1016/j.neuron.2020.01.026.

[102] Stefano Ambrogio, Pritish Narayanan, Hsinyu Tsai, Robert M Shelby, Irem Boybat, Carmelo Nolfo, Severin Sidler, Benjamin Killeen, Christina Cheng, Yassine Jaoudi, Massimo Giordano, Martina Bodini, and C P Nathan. “Equivalent-accuracy accelerated neural-network training using analogue memory”. In: Nature 558 (2018), pp. 60–67. issn: 1476-4687. DOI: 10.1038/s41586-018-0180-5. url: http://dx.doi.org/10.1038/s41586-018-0180-5.

[103] Chiara Bartolozzi, Giacomo Indiveri, and Elisa Donati. “Embodied neuromorphic intelligence”. In: Nature Communications 13.1 (2022), pp. 1–14. issn: 20411723. DOI: 10.1038/s41467-022-28487-2.

[104] Enming Song, Jinghua Li, Sang Min Won, Wubin Bai, and John A Rogers. “Materials for flexible bioelectronic systems as chronic neural interfaces”. In: Nature Materials 19.June (2020), pp. 590–603. issn: 1476-4660. DOI: 10.1038/s41563-020-0679-7. url: http://dx.doi.org/10.1038/s41563-020-0679-7.

[105] Ariel J Lee and Wenbo Wang. “Flexible and smart electronics for single-cell resolved brain – machine interfaces”. In: Applied Physics Review 10.011314 (2023), pp. 1–16. DOI: 10.1063/5.0115879. url: http://dx.doi.org/10.1063/5.0115879.

[106] Nicola Masala et al. “comorbidities of epilepsy Targeting aberrant dendritic integration to treat cognitive comorbidities of epilepsy”. In: Brain 146.6 (2023), pp. 2399–2417. issn: 0006-8950. DOI: 10.1093/brain/awac455. url: https://doi.org/10.1093/brain/awac455.

[107] Philip R Corlett, Guillermo Horga, Paul C Fletcher, Ben Alderson-day, Katharina Schmack, and Albert R Powers Iii. “Hallucinations and Strong Priors”. In: Trends in Cognitive Sciences 23.2 (2018), pp. 114–127. issn: 1364-6613. DOI: 10.1016/j.tics.2018.12.001. url: http://dx.doi.org/10.1016/j.tics.2018.12.001.

[108] Naoya Takahashi, Sara Moberg, Timothy A. Zolnik, Julien Catanese, Robert N.S. Sachdev, Matthew E. Larkum, and Dieter Jaeger. “Thalamic input to motor cortex facilitates goal-directed action initiation”. In: Current Biology 31.18 (2021), 4148–4155.e4. issn: 18790445. DOI: 10.1016/j.cub.2021.06.089. url: https://doi.org/10.1016/j.cub.2021.06.089.

[109] Huifang E Wang, Borana Dollomaja, Paul Triebkorn, Gian Marco Duma, Adam Williamson, Julia Makhalova, Jean-didier Lemarechal, Fabrice Bartolomei, and Viktor Jirsa. “Virtual brain twins for stimulation in epilepsy”. In: Nature Computational Science 5.September 2025 (2025), pp. 754–768. issn: 2662-8457. DOI: 10.1038/s43588-025-00841-6. url: http://dx.doi.org/10.1038/s43588-025-00841-6.

[110] Andrej Karpathy. tinyshakespeare dataset. https://github.com/karpathy/char-rnn/tree/master/data/tinyshakespeare. Accessed: 2026-08-03. 2015.

[111] Hugging Face Team. tokenizers: Fast and Easy Tokenizer Library. 2020. url: https://github.com/huggingface/tokenizers.

[112] Diederik P Kingma and Jimmy Ba. “Adam: A method for stochastic optimization”. In: arXiv preprint arXiv:1412.6980 (2014).

[113] Charles Kelly and Tatoeba Project. Tab-delimited Bilingual Sentence Pairs. http://www.manythings.org/anki/. Accessed: 2026-08-03.

[114] Richard S. Sutton and Andrew G. Barto. Reinforcement learning. 2nd. MIT Press, 2018. isbn: 9788578110796. arXiv: arXiv:1011.1669v3.

[115] Maja Popović. “chrF: character n-gram F-score for automatic MT evaluation”. In: Proceedings of the tenth workshop on statistical machine translation. 2015, pp. 392–395.

[116] Timothy Paul Lillicrap, Jonathan James Hunt, Alexander Pritzel, Nicolas Manfred Otto Heess, Tom Erez, Yuval Tassa, David Silver, and Daniel Pieter Wierstra. Continuous control with deep reinforcement learning. US Patent 10,776,692. 2020.

[117] Garofolo, John S., Lamel, Lori F., Fisher, William M., Pallett, David S., Dahlgren, Nancy L., Zue, Victor, and Fiscus, Jonathan G. TIMIT Acoustic-Phonetic Continuous Speech Corpus. 1993. DOI: 10.35111/17GK-BN40. (Visited on 01/05/2026).

[118] Nathan E Crone, Dana Boatman, Barry Gordon, and Lei Hao. “Induced Electrocorticographic Gamma Activity during Auditory Perception”. In: Clinical Neurophysiology 112.4 (Apr. 2001), pp. 565–582. issn: 13882457. DOI: 10.1016/S1388-2457(00)00545-9. (Visited on 01/18/2022).

[119] Yoav Benjamini and Yosef Hochberg. “Controlling the False Discovery Rate: A Practical and Powerful Approach to Multiple Testing”. In: Journal of the Royal Statistical Society 57.1 (1995), pp. 289–300.

## Supplementary References

[38] Virginie van Wassenhove, Ken W. Grant, and David Poeppel. “Temporal window of integration in auditory-visual speech perception”. In: Neuropsychologia 45.3 (2007), pp. 598–607. issn: 00283932. DOI: 10.1016/j.neuropsychologia.2006.01.001.

[120] C. Cappe, E. M. Rouiller, and P. Barone. “Multisensory anatomical pathways”. In: Hearing Research 258. 1-2 (2009), pp. 28–36. issn: 03785955. DOI: 10.1016/j.heares.2009.04.017. url: http://dx.doi.org/10.1016/j.heares.2009.04.017.

[121] Imanol Schlag, Kazuki Irie, and Jürgen Schmidhuber. “Linear transformers are secretly fast weight programmers”. In: International conference on machine learning. PMLR. 2021, pp. 9355–9366.

[122] Hao Peng, Nikolaos Pappas, Dani Yogatama, Roy Schwartz, Noah Smith, and Lingpeng Kong. “Random Feature Attention”. In: International Conference on Learning Representations. 2021.

[123] Krzysztof Marcin Choromanski, Valerii Likhosherstov, David Dohan, Xingyou Song, Andreea Gane, Tamas Sarlos, Peter Hawkins, Jared Quincy Davis, Afroz Mohiuddin, Lukasz Kaiser, David Benjamin Belanger, Lucy J Colwell, and Adrian Weller. “Rethinking Attention with Performers”. In: International Conference on Learning Representations. 2021.

[124] John Lisman, Ryohei Yasuda, and Sridhar Raghavachari. “Mechanisms of CaMKII action in long-term potentiation”. In: Nature Reviews Neuroscience 13.February (2012). issn: 1471-003X. DOI: 10.1038/nrn3192.

[125] Cristina M Alberini. “Transcription Factors in Long-Term Memory and Synaptic Plasticity”. In: Physiological Reviews 89 (2009), pp. 121–145. DOI: 10.1152/physrev.00017.2008.

[126] Yutao Sun, Li Dong, Shaohan Huang, Shuming Ma, Yuqing Xia, Jilong Xue, Jianyong Wang, and Furu Wei. “Retentive network: A successor to transformer for large language models”. In: arXiv preprint arXiv:2307.08621 (2023).

[127] Johannes Von Oswald, Eyvind Niklasson, Ettore Randazzo, João Sacramento, Alexander Mordvintsev, Andrey Zhmoginov, and Max Vladymyrov. “Transformers learn in-context by gradient descent”. In: International Conference on Machine Learning. PMLR. 2023, pp. 35151–35174.

[128] Wulfram Gerstner, Marco Lehmann, Vasiliki Liakoni, Dane Corneil, and Johanni Brea. “Eligibility traces and plasticity on behavioral time scales: experimental support of neoHebbian three-factor learning rules”. In: Frontiers in Neural Circuits 12 (2018), p. 53. DOI: 10.3389/fncir.2018.00053.

[129] Guillaume Bellec, Franz Scherr, Anand Subramoney, Elias Hajek, Darjan Salaj, Robert Legenstein, and Wolfgang Maass. “A solution to the learning dilemma for recurrent networks of spiking neurons”. In: Nature Communications 11 (2020), p. 3625. DOI: 10.1038/s41467-020-17236-y.

